# Exploring L-tyrosine and L-DOPA biosynthesis in faba bean (*Vicia faba* L.)

**DOI:** 10.64898/2026.02.26.707946

**Authors:** Xinxing Xia, Henryk Straube, Denise Blume, Davide Mancinotti, Bjørn Dueholm, Leandro Escobar-Herrera, Stig U. Andersen, Fernando Geu-Flores, Hester Sheehan

**Author notes:** Authors for correspondence: Hester Sheehan, Fernando Geu-Flores. Authors contributed equally (Xinxing Xia, Henryk Straube).

## Abstract

**Background and Aims:** L-DOPA is an important pharmaceutical that accumulates to high levels in the legume faba bean (*Vicia faba*). L-DOPA is likely derived from L-tyrosine but the responsible enzyme (L-tyrosine oxidase) remains unknown. Availability of L-tyrosine may be a key factor controlling L-DOPA accumulation. In legumes, L-tyrosine is supplied via either a plastidial TyrA enzyme (ADH) or a deregulated cytosolic homolog (PDH). This study aimed at identifying L-tyrosine oxidase and TyrA genes from faba bean.

**Methods:** We used gene-to-metabolite correlations and homology-based searches to select fifteen L-tyrosine oxidase candidates, which were tested in yeast and in the model plant *Nicotiana benthamiana*. We also used isotopically labeled L-tyrosine to measure biosynthetic activity in different faba bean tissues and to test an alternative biosynthetic hypothesis. Three faba bean TyrA genes were inferred by homology and assayed in *N. benthamiana* by co-expression with a known L-tyrosine oxidase, CYP76AD6.

**Key Results:** None of the L-tyrosine oxidase candidates produced L-DOPA upon heterologous expression. Feeding experiments showed a lack of correlation between L-DOPA accumulation and biosynthetic capacity. Feeding studies also disproved an alternative route to L-DOPA by oxidation of 4-hydroxyphenylpyruvate. Of the TyrA genes, two were able to increase L-tyrosine levels in *N. benthamiana* 2-3-fold (VfADH and VfPDH), and one of them was able to boost the levels of L-DOPA derivatives up to 6-fold (VfADH).

**Conclusions:** The faba bean L-tyrosine oxidase remains unidentified, with a possible transport of L-DOPA across tissues likely having confounded our correlation-based selection strategies. In *N. benthamiana*, both VfADH and VfPDH can increase the levels of L-tyrosine, while VfADH can further boost the levels of L-DOPA derivatives. Our work delivers a strategy to boost the provision of L-tyrosine in *N. benthamiana* and provides valuable insights in the search for the elusive L-tyrosine oxidase from faba bean.

## Introduction

L-DOPA (3,4-dihydroxyphenyl-L-alanine) is a non-proteinogenic amino acid derived from L-tyrosine with important ecological and defensive roles in plants. It accumulates to unusually high levels in two legume species (Fabaceae), faba bean (*Vicia faba*) and velvet bean (*Mucuna pruriens*; Damodaran and Ramaswamy 1937; Bell and Janzen 1971; Longo *et al*. 1974; Hill-Cottingham and Purves 1983; Duan *et al*. 2021), as well as in the latex of *Euphorbia* (Liß, 1961a, 1961b). L-DOPA acts as a potent allelochemical that suppresses competing vegetation and deters herbivores and pathogens (Rehr *et al*., 1973; Fujii *et al*., 1991; Nishihara *et al*., 2005; Soares *et al*., 2014). These effects are thought to arise from the rapid oxidation of L-DOPA to reactive quinones and melanin-like polymers that generate oxidative stress in surrounding organisms (Soares *et al*., 2014). L-DOPA acts furthermore as a below-ground pheromone involved in plant defense against herbivores (Cascone *et al*., 2023).

Besides its ecological importance, L-DOPA has high biomedical relevance as the main precursor of the neurotransmitter dopamine and as the gold standard for Parkinson’s disease treatment (Hornykiewicz, 2010). L-DOPA can cross the blood-brain barrier, after which it can be decarboxylated to give dopamine, thus compensating for dopamine loss in Parkinson’s patients. With cases expected to reach 12-17 million worldwide by 2040, reliable and affordable sources of L-DOPA are increasingly important (Dorsey *et al*., 2018; GBD 2016 Parkinson’s Disease Collaborators, 2018). At present, L-DOPA is mainly produced by chemical synthesis or microbial fermentation, relying on non-environmentally friendly reagents or addition of large amounts of precursors (Min *et al*., 2015; Jia *et al*., 2026). Instead, the use of plants or plant enzymes for L-DOPA production may offer a sustainable, cost-effective alternative.

The accumulation of L-DOPA in faba bean has been known since 1913 (Guggenheim, 1913). Feeding studies in the 1970s identified L-tyrosine as a precursor and excluded L-phenylalanine (Griffith and Conn, 1973), suggesting direct oxidation of L-tyrosine to give L-DOPA (Fig. 1A). Half a century later, however, the responsible enzyme remains unidentified. The only characterized L-tyrosine oxidases from plants are three closely related cytochrome P450s (CYPs), CYP76AD1/5/6, first characterized in beetroot (*Beta vulgaris*), and which are involved in pigment biosynthesis (DeLoache *et al*., 2015; Polturak *et al*., 2016; Sunnadeniya *et al*., 2016). All three can oxidize L-tyrosine to L-DOPA and CYP76AD1 additionally oxidizes L-DOPA to cyclo-DOPA. Both L-DOPA and cyclo-DOPA are converted further to betalain pigments (betaxanthin and betacyanin) with the aid of an unrelated oxidase, 4,5-DOPA dioxygenase (DODA; Fig. 1B; Sunnadeniya *et al*. 2016; Polturak *et al*. 2016). Heterologous expression of these oxidases in various plant systems has been shown to induce the accumulation of L-DOPA and betalain pigments (Polturak *et al*., 2017a; Breitel *et al*., 2020; Grützner *et al*., 2021).

**Figure 1.**
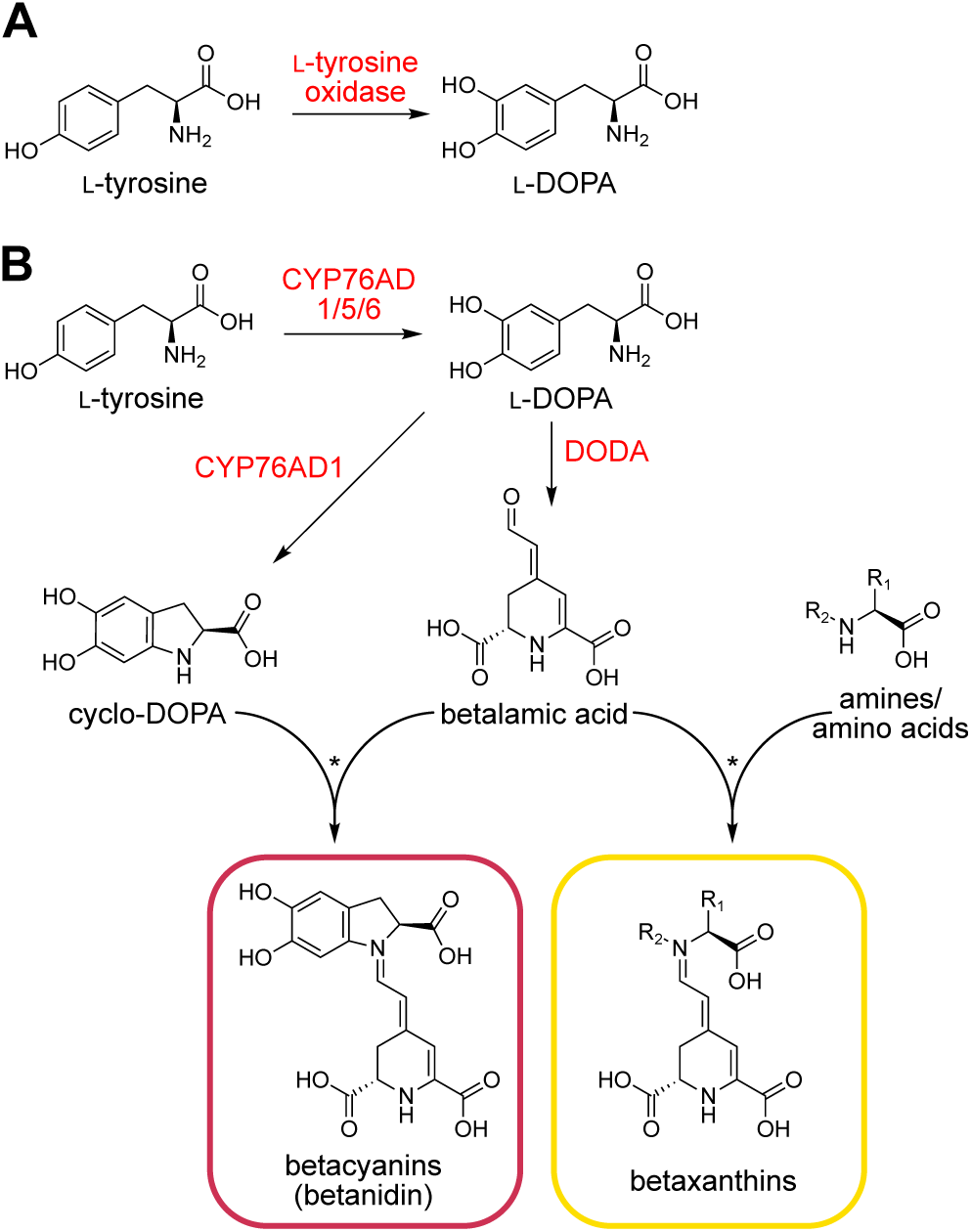
The proposed reaction for L-DOPA biosynthesis in faba bean, and the characterized reaction for L-DOPA biosynthesis leading to betalain production in beetroot. (A) In faba bean, L-tyrosine is converted to L-DOPA by an unknown L-tyrosine oxidase. (B) Enzymes from the CYP76AD subfamily convert L-tyrosine to L-DOPA in the first step of betalain biosynthesis in betalain-pigmented species (e.g. beetroot). DODA, L-DOPA 4,5-dioxygenase.

CYP76AD1/5/6 arose from gene duplications specific to the Caryophyllales, and no orthologs are expected in legumes (Brockington *et al*., 2015). Nevertheless, CYPs remain strong candidates for L-tyrosine oxidation because of their monooxygenase activity and central role in specialized metabolism (Hansen *et al*., 2021). Other plausible enzyme classes include 2-oxoglutarate/Fe(II)-dependent dioxygenases (2-ODD) and flavin monooxygenases (Mitchell and Weng, 2019). Polyphenol oxidases have also been implicated in L-tyrosine oxidation in velvet bean (Luthra and Singh, 2010; Araji *et al*., 2014; Nett *et al*., 2020; Saranya *et al*., 2021; Qiao *et al*., 2023) and in faba bean (Jena *et al*., 2024). However, early biochemical studies showed that although PPO activity of faba bean seedlings decreased when grown in the dark, L-DOPA levels remained unaffected (Griffith and Conn, 1973). This suggests that canonical PPOs are not involved in L-DOPA biosynthesis in faba bean.

The high accumulation of L-DOPA in faba bean is likely driven not only by the presence of an L-tyrosine oxidase but also by increased metabolic flux toward L-tyrosine. In plants, L-tyrosine is synthesized in plastids from prephenate via the consecutive action of prephenate aminotransferase (PPA-AT) and arogenate dehydrogenase (ADH). This plastidial pathway is tightly regulated by feedback inhibition of ADH by L-tyrosine (Gaines *et al*., 1982; Connelly and Conn, 1986; Rippert and Matringe, 2002a, 2002b). However, several plant lineages have evolved alternative biosynthesis pathways presumably to support production of tyrosine-derived specialized metabolites (Timoneda *et al*., 2018; Lopez-Nieves *et al*., 2022; El-Azaz *et al*., 2023, 2025). In Caryophyllales, ADH has undergone duplication, giving ADHα, an enzyme with reduced feedback sensitivity (Lopez-Nieves *et al*., 2018). In legumes, an additional cytosolic pathway arose, in which prephenate is first oxidized by prephenate dehydrogenase (PDH) and then transaminated to give L-tyrosine. PDH is evolutionarily related to ADH and is not inhibited by L-tyrosine (Rubin and Jensen, 1979; Schenck *et al*., 2017, 2020). PDH activity has been detected in faba bean extracts (Gamborg and Keeley, 1966; Schenck *et al*., 2020), but the corresponding genes have not been identified. Many legumes also contain a non-canonical ADH (ncADH) that shows only partial feedback inhibition (Schenck *et al*., 2017). Collectively, ADH and PDH enzymes are known as TyrA enzymes in reference to their bacterial counterparts (Schenck *et al*., 2017). While deregulated TyrA enzymes such as beet ADHα can increase the accumulation of L-tyrosine and downstream metabolites when expressed in *Nicotiana benthamiana* (Lopez-Nieves *et al*., 2018, 2022; Timoneda *et al*., 2018; Grützner *et al*., 2021; Jung and Maeda, 2024), the effects of legume PDH or ncADH variants have not been tested.

To shed light on the production of L-DOPA in faba bean, we aimed to identify the enzyme responsible for oxidizing L-tyrosine to L-DOPA as well as the TyrA enzymes involved in the provision of L-tyrosine. Among others, we used candidate selection approaches from a previous study where gene-to-metabolite correlations unveiled a key gene involved in the biosynthesis of the anti-nutrients vicine and convicine (Björnsdotter *et al*., 2021). We tested fifteen L-tyrosine oxidase candidates in two heterologous hosts, yeast (*Saccharomyces cerevisiae*) and *N. benthamiana*, but could not identify the enzyme. We then used isotopically labelled L-tyrosine to measure the biosynthetic capacity of different faba bean tissues and compared them to the endogenous L-DOPA contents. The weak correlation suggested L-DOPA mobility across tissues and provided an explanation for the lack of success of our correlation-based candidate selection strategies. We also used labeled precursor feeding to disprove an alternative hypothesis for L-DOPA biosynthesis by oxidation of 4-hydroxyphenylpyruvate (HPP). To investigate L-tyrosine biosynthesis, we identified three TyrA variants in faba bean (VfADH, VfncADH and VfPDH) and tested them in *N. benthamiana* by co-expression with CYP76AD6. Two of them increased the levels of L-tyrosine 2-3-fold (VfADH and VfPDH), and one of them further boosted the levels of L-DOPA derivatives up to 6-fold (VfADH). Our work delivers a strategy to improve L-tyrosine provision in *N. benthamiana* and provides valuable insights in the search for the elusive L-tyrosine oxidase from faba bean.

## Materials and methods

### Plant material and growth conditions

*V. faba* plants of the inbred line Hedin/2 and *N. benthamiana* plants (LAB strain; Ranawaka *et al*., 2023) were grown in the greenhouse under 16 h of light at 20 °C and 8 h of darkness at 19 °C at the Department of Plant and Environmental Sciences, University of Copenhagen (Frederiksberg, Denmark).

### Selection of candidate genes using gene-to-metabolite correlations

#### Correlation analysis via partial least squares regression

As a starting point, we used the transcriptomic and metabolomic datasets from Björnsdotter *et al*. (2021). Using the package edgeR (Robinson *et al*., 2009), we filtered both datasets by retaining metabolic features with an ion count > 20,000 in at least one sample and genes with a TPM value > 50 in at least one sample. The package mixOmics (Rohart *et al*., 2017; R Core Team, 2023) was used to carry out partial least squares (PLS) regression between the reduced datasets. The data were centered around zero and scaled to unit variance. Regression mode was used for the PLS deflation process, where the matrix X (genes and their expression levels) was used for the prediction of matrix Y (metabolic features and their ion counts). The correlations were calculated on the basis of individual samples for which we had both transcriptomics and metabolomics data. For the special case of whole seeds at early seed-filling-stage, the transcriptomics and metabolomics data had not been obtained from the same exact samples; thus, values across samples were averaged to give a single value per dataset. In total, the correlation analysis comprised 19 samples from the following tissues: young leaves (3), mature leaves (2), whole seeds at early seed-filling stage (1 averaged sample), pods at early seed-filling stage (3), pods at mid-maturation stage (2), embryos at mid-maturation stage (2), flowers (3), and stems (3).

Four metabolic features associated with L-DOPA [142_(+), 192_(+), 134_(+) and 120_(+)] were identified by comparing *m/z* values, retention times, and peak shapes to those obtained from the analysis of an L-DOPA standard using the same reversed-phase liquid chromatography coupled to mass-spectrometry (LC-MS) method as in Björnsdotter *et al* (2021; LC-MS analysis method A). A gene of interest, evgLocus_256923, was selected based on its annotation as a CYP and its close proximity to the location of the L-DOPA features within the correlation circle plot (Supplementary Fig. 1). To identify co-expressed genes, we used R to calculate Pearson correlation coefficients between genes using a larger transcriptomic dataset composed of all genes expressed above 20 TPM in at least one sample (Supplementary Data 1). The 50 most highly correlated genes were annotated according to the barrel clover (*Medicago truncatula)* reference genome (Tang *et al*., 2014) to find genes encoding putative oxidase enzymes.

#### Correlation analysis via Pearson correlation coefficients

We expanded the metabolomics dataset by Björnsdotter *et al*. (2021) with the addition of four extra tissues as well as a few additional samples of the original tissues. The new tissues/samples had been analyzed alongside the original ones (same LC-MS method and run) but had not been subjected to data analysis due to the lack of corresponding RNAseq data. The full metabolomics data comprises the following tissues, all derived from field-grown *V. faba* plants: young leaf (4 samples), mature leaves (3 samples), stem (4 samples), flower (4 samples), roots (4 samples), whole seeds at early seed-filling stage (3 samples), pods at early seed-filling stage (4 samples), seed coats at mid-maturation stage (1 sample), embryo at mid-maturation stage (3 samples), pods at mid-maturation stage (2 samples), pods at drying stage (4 samples), and seed coats at drying stage (4 samples). All samples were run in technical duplicates. We subjected the raw data to the same XCMS-based analysis pipeline described by Björnsdotter *et al*. (2021). Briefly, we used XCMS Online (v.3.7.1; Tautenhahn *et al*. 2012) to align chromatograms as well as identify and quantify metabolite features (peak width between 5 and 20 s; signal-to-noise ratio ≤ 6:1). We then removed metabolic features from the mass calibrant (retention time < 0.5 min) as well as those from blank samples (similar intensity between samples and blanks, *p* < 0.01 in Student’s t-test). The intensity of the remaining metabolic features was then normalized to the dry weight of the samples and to the signal of the internal standard ([M+H]^+^ for caffeine). Please note that the metabolic feature identifiers (IDs) in our expanded dataset will not necessarily correspond to those of the corresponding feature in the previously published dataset of Björnsdotter *et al*. (2021).

Prior to calculating correlation coefficients between genes and metabolites, the transcriptomics dataset from Björnsdotter *et al*. (2021) was reduced by removing genes with low variance (SD < 5). The cor function of R v.3.4.3 (R Core Team, 2023) was used to calculate the Pearson correlation coefficients for gene expression versus metabolite feature intensity. As with the PLS regression (see above), the correlations were calculated on the basis of individual samples except for whole seeds at early seed-filling-stage and, in total, the same 19 samples were analysed.

To identify the metabolite features associated with L-DOPA, we analyzed a commercial standard (Sigma-Aldrich, Germany) using the same LC-MS method (LC-MS method A below). By comparison of *m/z* ratios, retention times, and peak shapes, we selected four metabolic features corresponding to L-DOPA [42_(+), 143_(+), 107_(+), 41_(+)]. Genes were then ranked by their average correlation coefficients with respect to the four L-DOPA features. The top-250 genes were annotated, of which ten were found to encode putative oxidase enzymes. The short-list was further reduced to three candidates by examining the reactions that close homologues in other organisms were characterized to catalyze or predicted to do so and choosing those associated to reactions most chemically similar to the conversion of L-tyrosine to L-DOPA. The underlying data and code associated with the PCC analysis are available at TU Wien Research Data (doi: 10.48436/kq7h6-tgf87).

### Selection of candidate genes using homology to characterised enzymes

To select candidates homologous to CYP76AD6, 4-coumaroyl shikimate/quinate 3-hydroxylase (C3’H/CYP98A3), ascorbate peroxidase (APX), and TyrA enzymes, we used local BLASTp searches against the translated Hedin/2 transcriptome from Björnsdotter *et al*. (2021). For CYP76AD6 (NCBI accession AJD87474.1), we manually examined the top-50 local BLAST hits, searching for similarity of their expression profiles to the accumulation of L-DOPA as represented by metabolic features 142_(+), 192_(+), 134_(+) and 120_(+) from Björnsdotter *et al*. (2021) across eight tissues (young leaves, mature leaves, whole seeds at early seed-filling stage, pods at early seed-filling stage, pods at mid-maturation stage, embryos at mid-maturation stage, flowers, and stems). Two candidates were selected for testing: evgLocus_40272 and evgLocus_256923 (Supplementary Data 2). For C3’H/CYP98A3 (NCBI accession OAP09214.1), we selected the best BLAST hit, evgLocus_727341. For APX enzymes, we used AtAPX1 as query (NCBI accession AT1G07890.3) and examined the top-20 BLAST hits. APX candidates were selected further if they had a BLAST E-value lower than 2 x 10^-25^ (evgLocus_959121, evgLocus_1145931, evgLocus_145163, evgLocus_255817, evgLocus_189125, evgLocus_1209062) or an E-value lower than 2 x 10^-8^ and an expression level of at least 25 TPM in at least one sample (evgLocus_297956; Data S2). For TyrA enzymes, we used MtADH (Mt3g071980) and MtPDH1 (NCBI accession AIU94227.1) as BLASTp queries. The top three candidates (evgLocus_20583, evgLocus_220165, evgLocus_1024218; Data S2) were aligned to characterised legume TyrA protein sequences using the MAFFT plugin (Katoh and Standley, 2013) with default settings in Geneious Primer 2026.0.2 (https://www.geneious.com). To produce a phylogenetic tree, we supplemented the amino acid sequences from figure 1b of Schenck *et al*. (2017) with the faba bean TyrA candidates and MtADH and aligned the sequences with MAFFT v7.409 (Katoh & Standley, 2013) (--genafpair --maxiterate 1000). Using pxclsq from phyx (p 0.1; <u>Brown *et al*. 2017)</u>, the dataset was cleaned of columns with 90% or more missing data and a maximum likelihood tree was inferred using IQ-TREE (Nguyen *et al*., 2015) under the Whelan and Goldman model of sequence evolution with 1000 bootstrap replicates. Coding sequences (CDSs) of all L-tyrosine oxidase candidates are available in Supplementary Appendix 1 and faba bean TyrA genes have been deposited in Genbank (PZ024027-PZ024029).

### Cloning of candidate coding sequences for heterologous expression

Hedin/2 leaves were ground in liquid nitrogen and RNA was extracted using the Spectrum Plant Total RNA kit (Sigma-Aldrich, Germany) according to the manufacturer’s specifications. cDNA was synthesized using the iScript cDNA Synthesis Kit (Bio-Rad, Denmark) according to the manufacturer’s specifications. The cDNA was used to amplify the coding sequences shown in Appendix S1 by PCR using the primer pairs shown in Supplementary Table 1. The PCR products were cloned into heterologous expression vectors as described below.

For heterologous expression in *N. benthamiana*, oxidase candidates were cloned into pEAQ-USER (Luo *et al*., 2016) via USER cloning (Nour-Eldin *et al*., 2010) while TyrA candidates were cloned into pHREAC (Peyret *et al*. 2019; Addgene #134908) using either Golden Gate or Gibson Assembly (New England Biolabs, NEB; Chuang and Franke 2022). We also cloned the coding sequence of CYP76AD6 into pHREAC using Golden Gate Assembly starting from a plasmid template kindly provided by Adam Jozwiak and Asaph Aharoni (Weizmann Institute of Science, Israel).

Cloning of yeast expression vectors was carried out using Golden Gate Assembly and the yeast toolkit designed by Lee *et al*. (2015). *Beta vulgaris* DODAα1 (MN153194) was cloned from the CDS part plasmid pL0-BvDODAα1 (Guerrero-Rubio *et al*., 2023) into the URA3 integration vector pYTK096, along with the strong yeast promoter pTDH3 (pYTK009) and terminator tTDH1 (pYTK056). The yeast expression cassette plasmid pEVY002 was assembled from parts in the toolkit including the strong constitutive yeast promoter pCCW12 (pYTK010), an mRFP expression dropout for red/white screening (pYTK001), the yeast terminator tADH1 (pYTK053), the LEU2 locus (pYTK075), the CEN6/ARS4 replication locus (pYTK081), and ampicillin resistance (pYTK083). Subsequently, an empty pEVY002 vector without the RFP transcriptional unit was assembled by amplifying the backbone with Q5 High-Fidelity polymerase (NEB, USA), phosphorylating the 5’ ends of the purified PCR product with T4 polynucleotide kinase (Promega, USA), and ligating with T4 DNA ligase (NEB, USA). The CDSs of candidate genes were amplified from the pEAQ-USER constructs described above using Q5 High-Fidelity polymerase (NEB, USA) and cloned into pEVY002 (see primer pairs in Table S1). CYP76AD and CYP76AD6 were cloned from the CDS part plasmids by Guerrero-Rubio *et al*. (2023). One sequence (evgLocus_145163) was cloned using NEBuilder® HiFi DNA Assembly (NEB, USA) according to the manufacturer’s specifications.

### Heterologous expression of candidate genes in yeast

The *Saccharomyces cerevisiae* strain yWCD230 ([BY4741]his3Δ0, kindly provided by J. E. Dueber, University of California, USA) was transformed with BvDODAα1-pYTK096 following linearization with NotI, producing the strain ScDODAa1. The pEVY002 constructs carrying candidate genes were transformed into ScDODAa1, and the pEVY002 empty plasmid was also transformed to produce a negative control strain. Transformations were carried out using the *S.c.* EasyComp™ Transformation Kit (Thermo Fisher Scientific, USA) and strains were confirmed using colony PCR. Yeast strains were cultured in synthetic defined (SD) media under auxotrophic selection (SD-Ura for yHS-DODAa1; SD-Ura/-Leu for strains carrying pEVY002). The assay was carried out as described by Guerrero-Rubio *et al* (2023). In brief, strains were pre-cultured for 48 h in 3 mL of SD-Ura/-Leu at 30 °C at 200 rpm. Ten μL of saturated cultures were diluted into 500 μL of fresh media supplemented with 1 mM L-tyrosine and grown in a 96 deep-well plate at 30 °C at 300 rpm. Each strain was tested in four replicates. After 24 h, 100 μL of cells were pelleted by centrifugation in a 96-well PCR plate, rinsed in phosphate-buffered saline (PBS) pH 7.4, resuspended in 125 μL of PBS, and 100 μL was transferred to a black-walled, clear-bottomed 96-well plate. Cell density and intracellular fluorescence were quantified using a GloMax®-Multi Detection System (Promega, USA) using the Blue Optical Kit (excitation 490 nm/ emission 510-570 nm). Values were normalized based on the fluorescence of the negative control strain and corrected by the cell density (OD600).

### Transient expression of candidate genes in *N. benthamiana*

Transient expression in *N. benthamiana* was performed according to Chuang & Franke (2022) with some modifications. All plasmids were transformed into the *Agrobacterium tumefaciens* strain AGL-1 and grown in YEP media containing the antibiotics carbenicillin (50 µg mL^-1^), rifampicin (50 µg mL^-1^) and kanamycin (50 µg mL^-1^). Transformants were confirmed by colony PCR. A single positive colony was grown at 28 °C overnight in 5 ml of liquid YEP media with the same antibiotics. The cultures were maintained at 28 °C and 220 rpm until they reached an OD600 of 1. The cells were then harvested by centrifugation and resuspended in MMA buffer [10 mM MgCl2, 10 mM MES [2-(*N*-morpholino)ethanesulfonic acid], and 100 µM acetosyringone, pH 5.6] to give a final OD600 of 0.4-0.6. The resuspensions were used to infiltrate the third true leaf of 3-4-week-old plants in 3 replicates (each replicate corresponding to a different plant). To test TyrA genes, strains carrying the CDSs in the pHREAC plasmid were mixed with an analogous strain carrying GFP (kindly provided by Hadrien Peyret and George Lomonossoff, John Innes Centre, United Kingdom) or CYP76AD6 in equal volumes before infiltration. The plants were returned to the greenhouse following infiltration. After five days, infiltrated areas were collected as three 0.4-cm leaf discs and stored at −80 °C for later analysis. Metabolites were extracted and analyzed according to LC-MS analysis method B.

### Feeding labelled L-tyrosine to faba bean tissues

L-tyrosine-^13^C9 and L-tyrosine-^13^C2D5^15^N were purchased from Cambridge Isotope Laboratories (USA). Seeds of the Hedin/2 inbred line were germinated and grown in the greenhouse (see above for growth conditions). A volume of 1.5 ml of labelled compound was fed to different tissues at a concentration of 1 mM for 24 h at room temperature under constant light conditions. As controls, unlabeled L-tyrosine was fed in an identical way. For the feeding of both labelled compounds to radicles, we used five-day-old germinating seeds. For the feeding of all other tissue types with L-tyrosine-^13^C9, we used two-week-old seedlings. The latter tissue types were fed in different ways as follows. For leaves, we fed isolated leaves through the petiole. For seedling roots, whole seedlings were fed through the roots. For stems, internodal stem segments carrying a leaf were fed via the cut stem. Incorporation of L-tyrosine-^13^C9 was analyzed using metabolite extraction and LC-MS analysis method A (see below). Incorporation of L-tyrosine-^13^C2D5^15^N was analyzed by metabolite extraction and LC-MS analysis method B (see below).

### Metabolite extraction and LC-MS analysis method A

Tissues were ground in liquid nitrogen and 50-100 mg per sample were extracted in 250 μL extraction buffer A (60% methanol, 0.06% formic acid, and 5 ppm caffeine as internal standard). The mixtures were vortexed for 2 min and incubated at room temperature for 2 h with shaking at 1,200 rpm. After centrifugation for 10 min at 20,000 g at room temperature, 40 μL of the supernatant was diluted into 160 μL H2O. The samples were filtered through a 0.22-μM PVDF filter and transferred to HPLC vials with a glass insert. Metabolite analysis was carried out using LC-MS as described by Björnsdotter *et al*. (2021), including normalization by sample weight and internal standard. The L-DOPA peak ([M+H]^+^) was identified by comparison to a commercial standard.

### Metabolite extraction and LC-MS analysis method B

We tested L-tyrosine oxidase candidates and TyrA candidates in *N. benthamiana* using an optimized LC-MS method that gave improved peak shape for L-DOPA compared to the method by Björnsdotter *et al*. (2021). We also used the optimized method for feeding experiments with L-tyrosine-^13^C2D5^15^N.

Leaf and root tissues were extracted slightly differently. Leaf discs were ground in a mixer mill (Retsch, Germany) at 30 hz for 1 min using 1.5 ml safe-lock reaction tubes containing 3 steel-balls per tube. A volume of 250 μL extraction buffer B (60% methanol and 0.1% formic acid, as well as 50 µM caffeine and 50 μM uridine as internal standards) was added and samples were extracted in an overhead shaker for 1 h at 4°C. The samples were centrifuged for 10 min at 21,000 g at room temperature, and 40 μL of the supernatant was diluted into 160 μL H2O. Root tissues were ground in a similar way compared to leaf discs, but for 3 min and using five steel balls. A volume of 500 µl extraction buffer B was added, and samples were extracted at room temperature for 30 min at 800 rpm. The samples were then centrifuged for 10 min at 21,000 g at room temperature, and 20 μL of the supernatant was diluted into 180 μL H2O.

All samples were filtered through a 0.22 μM PVDF filter and transferred to HPLC vials with a glass insert. The analysis was carried out using the same LC system, qTOF detector, and source parameters as in LC-MS analysis method A above. A recently published chromatography method from Leite Dias *et al*. (2023) was adapted as follows. Separation was performed on a Waters Acquity UPLC HSS T3 column (2.1 mm × 100 mm, 1.8 µm) equipped with a Acquity UPLC HSS T3 VanGuard pre-column (1.8 µm, 2.1 × 5 mm; Waters, Germany) at 30 °C and 0.4 mL min^−1^ of flow rate. Mobile phases A and B consisted of 0.05% formic acid in water and 0.05% formic acid in acetonitrile, respectively. The following elution profile was used: 0–3 min, 0.1% B (constant); 3–6 min, 0.1–5% B (linear); 6–10 min, 5–12.5% B (linear), 10–10.5 min, 12.5–50% B (linear); 10.5–11 min, 50% B (constant); 11–11.5 min, 50– 99% B (linear); 11.5-13 min, 99% B (constant), 13-13.5 min 99-0.1%, and 13.5–16 min, 0.1% B (constant). L-DOPA and dopamine peaks were confirmed by comparison to commercial standards with respect to *m/z* ratios, fragmentation pattern, peak shape, and retention time.

### Statistical analysis

For analysis of fluorescence levels (Fig. 4A) and metabolites, including isotopically labeled analytes (Figs. 5, 6, 8 and S3) statistical significance was assessed using GraphPad Prism 10 (www.graphpad.com), including the calculation of standard deviation shown in the graphs as described in figure legends. The number of replicates is also reported in the figure legends. Supplementary Data 3 provides the raw datasets underlying the graphs in the relevant figures and supplementary figures.

## Results

### Analysis of L-DOPA in eight faba bean tissues

In a previous study, we had generated transcriptomic and metabolomic datasets for eight aerial tissues of the field-grown, inbred faba bean line Hedin/2 (Björnsdotter *et al*., 2021). In the present study, we mined the metabolomic dataset for metabolic features corresponding to L-DOPA by comparison to a commercial standard that ran with the same LC-MS method. The selected features corresponded to the protonated form of L-DOPA [feature 142_(+)] as well as three in-source fragments arising from decarboxylation [feature 192_(+)], deamination [feature 134_(+)], and loss of the amino acid moiety [feature 120_(+)]. All three fragments were readily identifiable in MS2 spectra of protonated L-DOPA (Fig. 2A). Analysis of these features across tissues revealed that L-DOPA accumulated the most in flowers, pods at early seed-filling and mid-maturation stages, and whole seeds at early seed-filing stages, while stems, young leaves, and mature leaves accumulated lower levels (Fig. 2B). Only traces of L-DOPA were observed in embryos at mid-maturation stage (Fig. 2B), which is consistent with previous studies (Longo *et al*., 1974; Hill-Cottingham & Purves, 1983; Duan *et al*., 2021).

**Figure 2.**
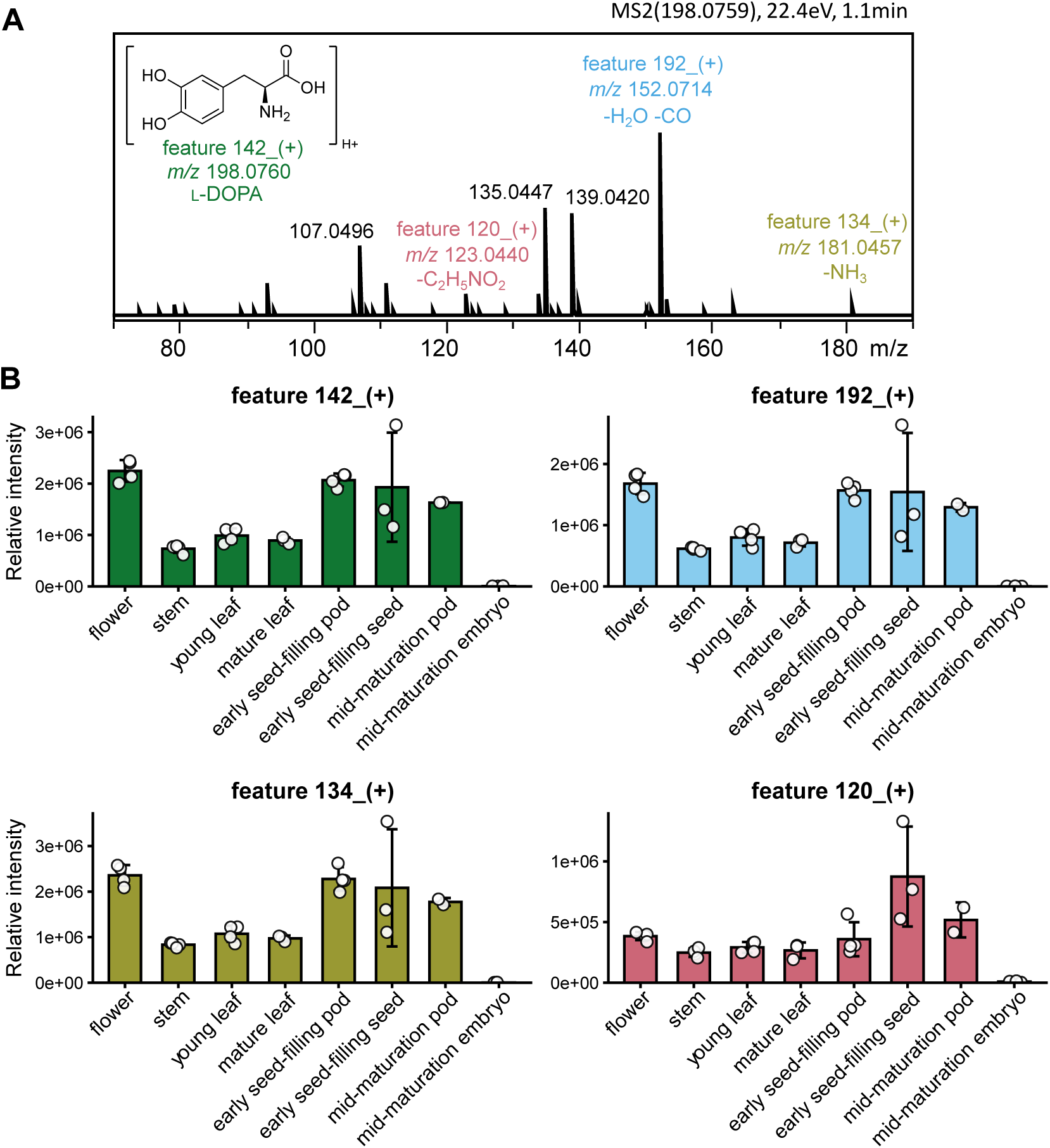
Metabolic features representing L-DOPA, and their abundance in eight aerial faba bean tissues as inferred from the metabolomic dataset by Björnsdotter *et al*. (2021). (A) MS2 spectra of a commercial standard of L-DOPA. The selected metabolic features are indicated above the corresponding daughter ion signals (colored labels). (B) Relative levels of L-DOPA in different tissues of faba bean as indicated by the mean abundance of L-DOPA-associated metabolic features across faba bean tissues. The intensities of metabolic features were normalized to the internal standard (caffeine) and to the dry weight of each sample. Hollow circles represent individual data points, bars represent mean values, and error bars represent the standard deviation (±).

### Selection of faba bean candidate genes encoding L-tyrosine oxidase

The most common method to uncover enzymes in plant specialized metabolism relies on gene co-expression, where a known biosynthetic gene is used as bait to discover others in the same pathway (Delli-Ponti *et al*., 2021). However, the pathway to L-DOPA is likely a single-enzyme pathway, rendering this method unapplicable. An alternative method uses both RNA-seq and metabolite data to implicate genes whose expression is correlated with accumulation of the metabolite of interest across tissues (Polturak *et al*., 2017b; Tu *et al*., 2020; Colinas *et al*., 2025). Candidates are then prioritized according to the class of enzyme that they may encode, considering the likelihood of catalyzing the target reaction. Here, we selected candidate genes coding for L-tyrosine oxidase genes by carrying out gene-to-metabolite correlation analyses with the Björnsdotter *et al*. (2021) datasets and by homology to characterized enzymes that carry out similar reactions in other species.

Our first gene-to-metabolite correlation analysis was based on partial least squares regression using the R package mixOmics (see datasets and script in Data S1; Rohart *et al*. 2017; R Core Team 2023). This generated a correlation circle plot where the proximity between genes and metabolites represents a degree of positive correlation (Fig. S1; González *et al*. 2012). A gene predicted to encode a CYP736A21 enzyme, evgLocus_256923, was found close to a cluster of the L-DOPA associated metabolite features 142_(+), 192_(+), and 134_(+), and was therefore inferred to be correlated with L-DOPA accumulation (Table 1; Fig. S1). We then used this gene as a bait in a co-expression analysis on the same transcriptomic dataset and selected a homolog of it, evgLocus_1282817, with a Pearson correlation coefficient of 0.86 (Table 1; Data S1).

**Table 1.** Gene candidates for L-tyrosine oxidase activity in faba bean. 2-ODD, 2-oxoglutarate-dependent dioxygenase; APX, ascorbate peroxidase; C3’H, 4-coumaroyl shikimate/quinate 3’-hydroxylase; CYP, cytochrome P450; PCC, Pearson correlation coefficient analysis; PLS, partial least squares regression analysis.

| Gene ID | Implicated by | Class of enzyme | Closest homologue in barrel clover |
| --- | --- | --- | --- |
| evgLocus_1242091 | PCC | CYP | CYP720B1 |
| evgLocus_30215 | PCC | CYP | CYP714A1 |
| evgLocus_1273529 | PCC | 2-ODD | 2'-deoxymugineic-acid 2'-dioxygenase-like protein |
| evgLocus_1282817 | PLS | CYP | CYP71A21 |
| evgLocus_256923 | PLS | CYP | CYP71A21 |
| evgLocus_40272 | CYP76AD6 homologue | CYP | CYP71A21 |
| evgLocus_366734 | CYP76AD6 homologue | CYP | CYP76C4 |
| evgLocus_959121 | APX homologue | APX | L-ascorbate peroxidase |
| evgLocus_89125 | APX homologue | APX | L-ascorbate peroxidase 6 |
| evgLocus_145163 | APX homologue | APX | L-ascorbate peroxidase |
| evgLocus_1209062 | APX homologue | APX | putative L-ascorbate peroxidase 6 |
| evgLocus_145931 | APX homologue | APX | L-ascorbate peroxidase |
| evgLocus_255817 | APX homologue | APX | L-ascorbate peroxidase |
| evgLocus_297956 | APX homologue | APX | L-ascorbate peroxidase |
| evgLocus_727341 | C3'H homologue | CYP | flavonoid 3'-monooxygenase |

We then carried out a separate gene-to-metabolite analyses solely based on Pearson correlation. For this, we first updated the metabolomics dataset by Björnsdotter *et al*. (2021) to include a few additional tissues that had been harvested and analyzed alongside the original ones but had not been subject to data analysis (roots, seed coats at mid-maturation stage, pods at drying stage, and seed coats at drying stage). In this updated dataset, the L-DOPA features mentioned above were assigned different identifiers, namely 42_(+), 143_(+), 107_(+), and 41_(+) (Supplementary Fig. 2). The transcriptomic dataset was updated to filter out genes that showed little difference across samples and we subsequently calculated Pearson correlation coefficients between all genes and metabolites. Focusing on the average correlation coefficients to the four L-DOPA features, we mined the top 250 ranked genes for oxidative enzymes and obtained two candidates predicted to encode CYP enzymes, evgLocus_1242091 (average PCC of 0.72; CYP720B1) and evgLocus_30215 (average PCC of 0.60; CYP714A1), as well as one candidate predicted to encode a 2-ODD enzyme, evgLocus_1273529 (average PCC of 0.66; 2’-deoxymugineic-acid 2’-dioxygenase-like; Table 1).

Our remaining gene candidates were selected by homology to enzymes carrying out L-tyrosine hydroxylation or similar reactions. We first mined plant metabolism for similar reactions, hydroxylating a phenolic ring at the *ortho* position and focused on two such instances. First, an APX from thale cress (*Arabidopsis thaliana*) was shown to catalyze the conversion of *para*-coumaric acid to caffeic acid (Fig. 3A; Barros *et al*., 2019). Interestingly, the recombinant APX enzyme can catalyze the conversion of L-tyrosine to L-DOPA *in vitro* (Barros *et al*., 2019). Second, the enzyme C3’H (CYP98A3) involved in the lignin biosynthesis pathway catalyzes the hydroxylation of coumaroyl shikimate to caffeoyl shikimate (Fig. 3B; Schoch *et al*., 2001). With respect to APXs, we used local BLAST searches against the Hedin/2 transcriptome to select seven candidate genes with similarity to the mentioned thale cress APX: evgLocus_959121, evgLocus_89125, evgLocus_145163, evgLocus_1209062, evgLocus_145931, evgLocus_255817, and evgLocus_297956 (Table 1; Data S2). With respect to C3’H, we selected the top BLASTp hit when using thale cress C3’H (Schoch *et al*., 2001) as a query, namely evgLocus_727341 (79.8% identity; Table 1).

**Figure 3.**
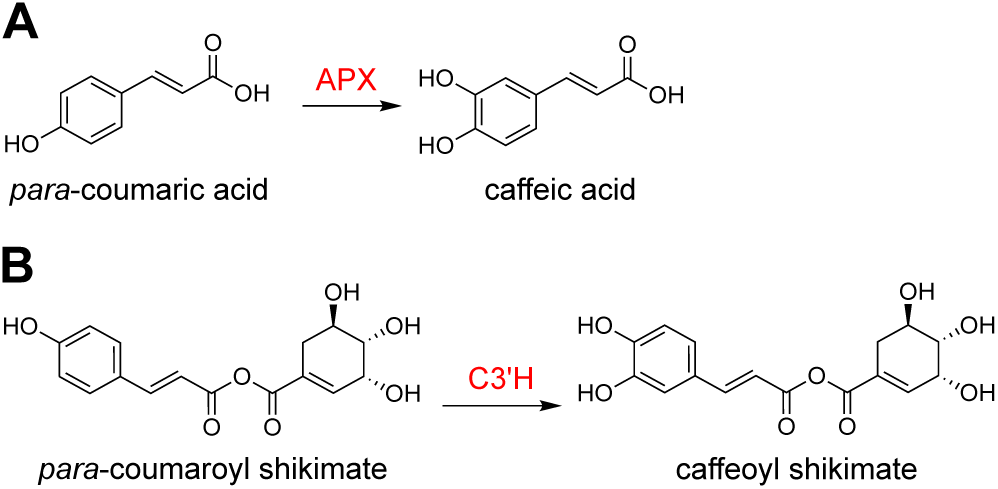
Chemical reactions in other plant systems that are analogous to the biosynthesis of L-DOPA. (A) An ascorbate peroxidase (APX) converts *para*-coumaric acid to caffeic acid in thale cress. (B) A cytochrome P450, 4-coumaroyl shikimate/quinate 3-hydroxylase (C3’H/CYP98A3), catalyzes the production of caffeoyl shikimate in thale cress.

CYP76AD6 hydroxylates tyrosine to L-DOPA in beetroot (Polturak *et al*., 2016; Sunnadeniya *et al*., 2016). Although we did not expect to find close homologs in faba bean (see Introduction), we searched for candidate genes among the top 50 BLASTp hits against the Hedin/2 transcriptome. We eliminated lowly expressed genes and then visually examined the expression patterns across tissues, selecting evgLocus_40272 (CYP71A21; 34.2% identity) and evgLocus_366734 (CYP76C4; 32.8% identity) as the only additional candidates (Table 1; Data S2). We also noted that three other CYP candidates implicated by other analyses were also present in this top 50 list: evgLocus_256923, evgLocus_1282817 and evgLocus_727341 (see above).

### Testing of candidate genes in yeast and *Nicotiana benthamiana*

To initially test our candidate genes, we used a yeast biosensor in which the production of the fluorescent yellow betalain pigments, betaxanthins, are a reporter for L-tyrosine hydroxylation activity (DeLoache *et al*., 2015; Savitskaya *et al*., 2019). We constructed the yeast strain ScDODAa1, which contains a chromosomally integrated *B. vulgaris* DODAα1 gene that enables the strain to convert L-DOPA to betalamic acid, which spontaneously condenses with free amines to form betaxanthins (Fig. 1B). If a strain makes betaxanthins when it expresses a candidate gene, that gene must encode an enzyme that converts L-tyrosine to L-DOPA. We expressed our candidates in ScDODAa1 under control of a strong yeast promoter, grew the cultures for 24 h with supplementation of 1 mM L-tyrosine, and measured their fluorescence. Two positive control strains that had been transformed with the tyrosine hydroxylation genes from beetroot, CYP76AD5 and CYP76AD6, produced strains that fluoresced and were noticeably orange (Fig. 4A). However, none of the strains expressing our L-tyrosine oxidase candidates produced fluorescence above the level of the negative control strain (Fig. 4A).

**Figure 4.**
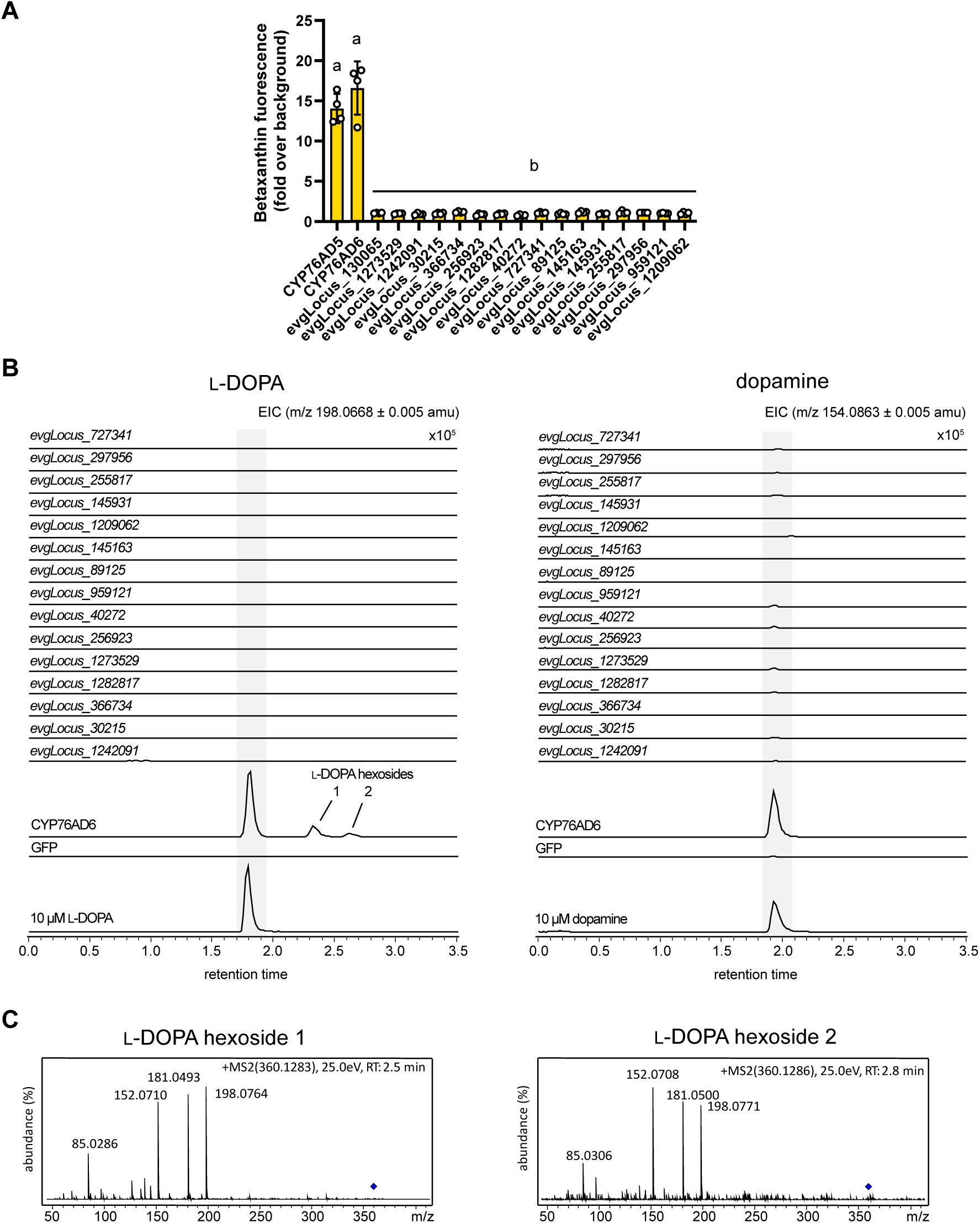
Screening candidates for L-tyrosine oxidase activity. (A) Quantification of betaxanthin fluorescence in yeast strains expressing BvDODAα1 and individual gene candidates, showing that none of the candidates led to betaxanthin production. CYP76AD5 and CYP76AD6 from beetroot were used as positive controls for L-tyrosine oxidase activity. Values were normalized to a strain expressing only BvDODAα1. Hollow circles represent individual data points, bars represent mean values, and error bars represent the standard deviation (±). Letters indicate significant differences in the mean values, identified by two-way ANOVA and Tukey’s post-hoc tests (*p* < 0.05). (B) Representative extracted ion chromatograms for L-DOPA and dopamine detected in leaves of *N. benthamiana* expressing the individual gene candidates. CYP76AD6 from beet root was used as positive control, and GFP was used as a negative one. L-DOPA and L-DOPA hexosides were only detected in the positive control. Dopamine was detected in high levels in the positive control and in trace amounts for the GFP control and for some of the candidates. For comparison, the equivalent chromatograms from commercial standards of L-DOPA and dopamine are shown. Chromatogram traces are shown for a single biological replicate; altogether three replicates were analyzed, all showing a similar result. (**C**) MS2 spectra of the L-DOPA hexosides 1 and 2 detected in *N. benthamiana* leaves expressing CYP76AD6 (naming corresponds to the peaks shown in panel B).

Considering that we did not detect L-tyrosine oxidase activity from any of our candidates in a microorganismal host, we decided to test the candidates in a plant host to more faithfully replicate their native cellular environment. We expressed the candidates transiently in leaves of *N. benthamiana* using agroinfiltration and included CYP76AD6 as a positive control as well as GFP as a negative control. After five days, we collected leaf discs and analyzed metabolites in methanolic extracts by LC-MS. We confirmed that overexpression of beetroot CYP76AD6 led to accumulation of L-DOPA, but L-DOPA could not be detected in any of the leaf samples from overexpression of our candidate genes (Fig. 4B). Interestingly, not only did we detect L-DOPA in the CYP76AD6 sample but also two additional compounds, as well as dopamine (Fig. 4B). Examination of MS and MS2 spectra suggested that the two unknown compounds were L-DOPA hexosides, likely glycosylated at either the 3’ or 4’-OH group (*m/z* around 360.1284 with familiar fragments in the MS2 spectra; Fig. 4C).

### Investigating the sites of biosynthesis of L-DOPA in faba bean

As long-distance L-DOPA transport may have had an impact on our correlation-based gene discovery strategies (see Discussion), we performed a precursor feeding experiment to identify the sites of L-DOPA biosynthesis. As it has been confirmed that L-tyrosine is a precursor of L-DOPA in faba bean (Griffith and Conn, 1973), we fed L-tyrosine-^13^C9 (Fig. 5A) to four different tissues: the radicle, seedling stems, seedling leaves, and seedling roots. The feeding was carried out for 24 hours under constant light at room temperature (Griffith and Conn, 1973). As a negative control, we carried out feeding of the same tissues using unlabeled L-tyrosine. Upon analysis of L-DOPA-^13^C9 using LC-MS, we observed L-DOPA-^13^C9 accumulation in all four tissues, with the highest accumulation in radicles (Fig. 5B). L-DOPA-^13^C9 was not detected in the four tissues that were fed with unlabeled L-tyrosine (Fig. 5B). We also analyzed unlabeled L-DOPA in the same LC-MS runs and found that it accumulated to a larger degree in seedling leaves and seedling stems compared to the radicle (Fig. 5C). The discrepancy in biosynthetic activity (highest in radicles) versus accumulation levels (highest in seedling leaves and stems) opens the possibility that long-distance transport plays a role in the distribution of L-DOPA in faba bean.

**Figure 5.**
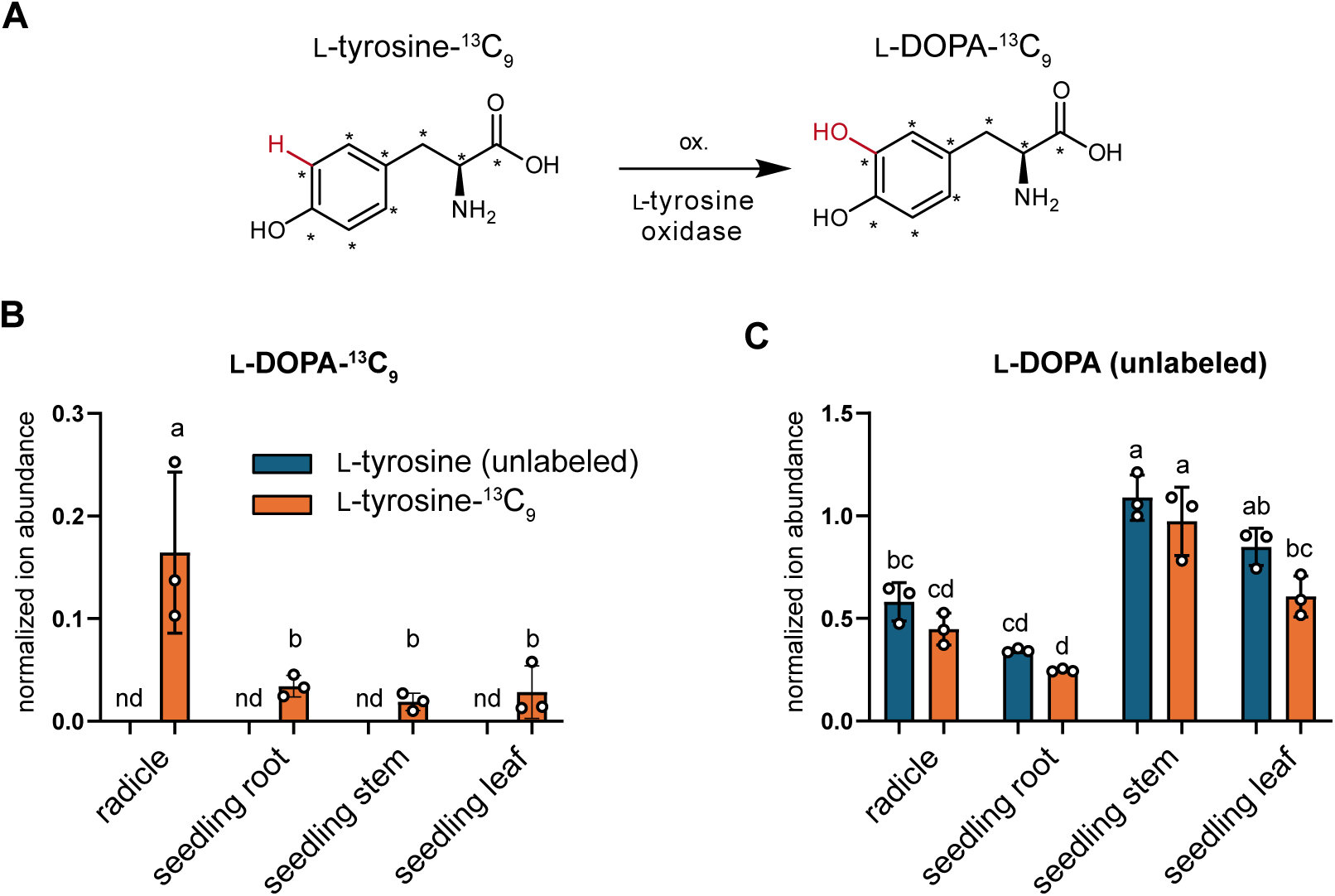
Precursor feeding experiments in seedling tissues of faba bean using L-tyrosine-^13^C9. (A) Expected conversion of L-tyrosine ^13^C9 to L-DOPA-^13^C9. (B) LC-MS quantification of L-DOPA-^13^C9 and unlabeled L-DOPA from radicles, seedling stems, seedling leaves, and seedling roots that were fed with either L-tyrosine-^13^C9 (orange bars) or unlabeled L-tyrosine (blue bars). Quantification was carried out relative to the internal standard, caffeine. Hollow circles represent individual data points, bars represent mean values, and error bars represent the standard deviation (±). Letters indicate significant differences in the mean values, identified by two-way ANOVA and Tukey’s post-hoc tests (*p* < 0.05). nd, not detected.

### Testing an alternative hypothesis for L-DOPA biosynthesis in faba bean

Based on the observed conversion of labelled L-tyrosine to L-DOPA in faba bean tissues (see above as well as the Introduction section), L-DOPA is thought to be derived from L-tyrosine by direct oxidation. We decided to test an alternative hypothesis that is consistent with the published feeding experiments. This hypothesis was inspired by the fact that the last step in the cytosolic PDH route to L-tyrosine biosynthesis in plants is reversible, namely, the transamination of HPP into L-tyrosine (Fig. 6A; Maeda and Dudareva 2012; Wang *et al*. 2016). Due to this reversibility, any L-tyrosine in faba bean is expected to be in equilibrium with HPP. We hypothesized that the oxidase of interest may hydroxylate HPP to give 3,4-dihydroxyphenylpyruvate (3-OH-HPP). The resulting 3-OH-HPP would then just need to be transaminated to give L-DOPA (Fig. 6A). To test this hypothesis, we obtained L-tyrosine-^13^C2D5^15^N (labelled at the amino nitrogen) and fed it to faba bean radicles for 24 hours. Unlabeled L-tyrosine was fed to the radicle as a negative control (Supplementary Fig. 3). If L-DOPA was derived from this alternative route, then the ^15^N would be lost during the reverse-transamination of L-tyrosine-^13^C2D5^15^N to HPP-^13^C2D5, ultimately giving L-DOPA-^13^C2D5. We detected L-tyrosine-^13^C2D5^15^N in fed radicles, indicating uptake of the labelled precursor, and we measured accumulation of L-DOPA-^13^C2D5^15^N in them, but could only detect trace amounts of L-DOPA-^13^C2D5 (Fig. 6B). This indicates that L-DOPA is not derived from this alternative biosynthetic pathway in faba bean.

**Figure 6.**
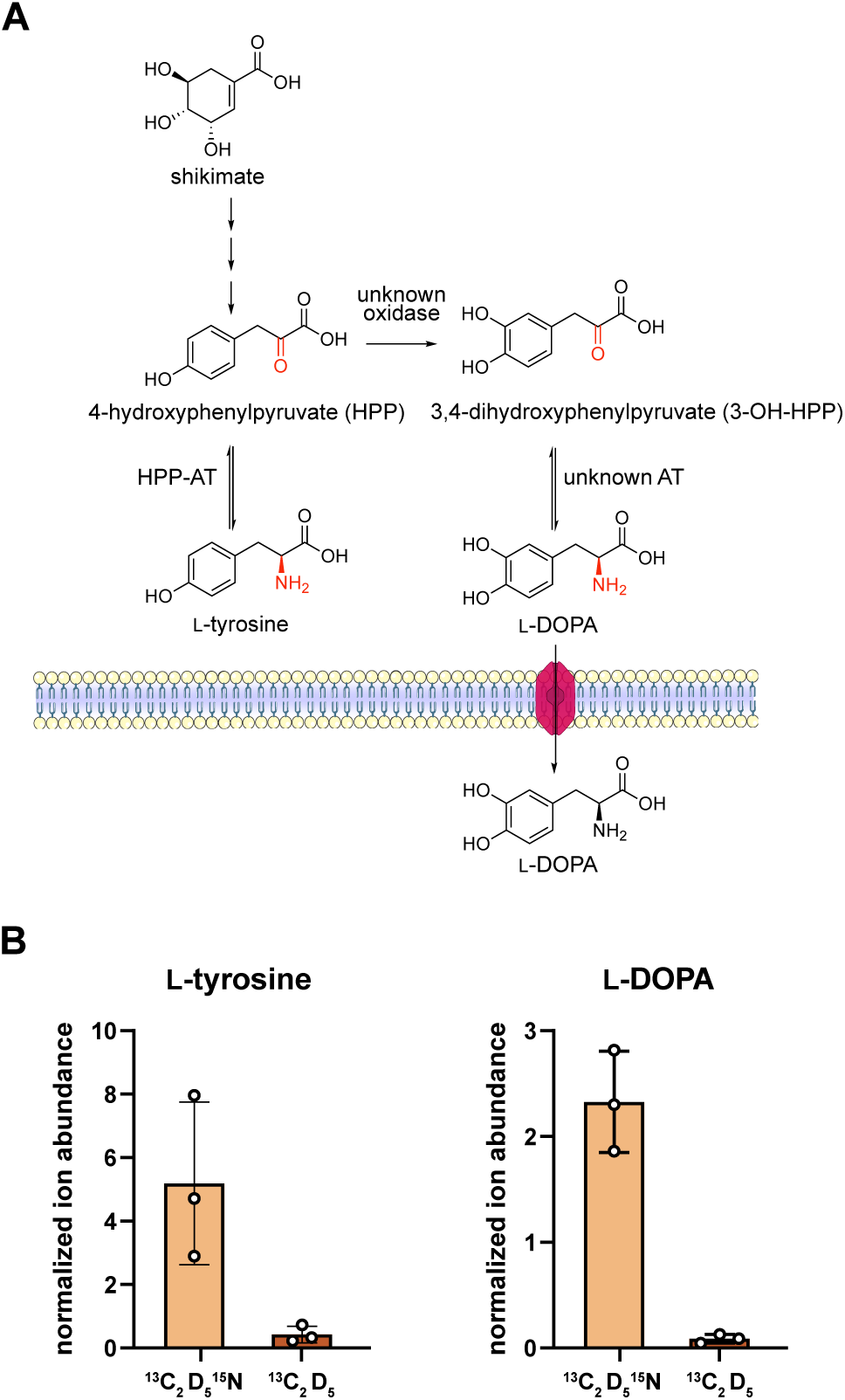
Testing 3,4-dihydroxyphenylpyruvate (3-OH-HPP) as an alternative precursor for L-DOPA. (A) The proposed reaction pathway in which L-DOPA is produced via transamination of 3-OH-HPP, analogous to the production of L-tyrosine from 4-hydroxyphenylpyruvate (HPP) via a reversible transamination reaction. This would require oxidation of HPP to 3-OH-HPP and transamination of 3-OH-HPP, as well as transport of L-DOPA away from the hypothesized transamination enzyme. (B) LC-MS quantification of ^13^C2D5^15^N (light orange) or ^13^C2D5 (dark orange) labeled L-tyrosine and L-DOPA in faba bean seedlings that were fed L-tyrosine-^13^C2D5^15^N. Quantification was carried out relative to the internal standard, caffeine. Hollow circles represent individual data points, bars represent mean values, and error bars represent the standard deviation (±). HPP-AT, 4-hydroxyphenylpyruvate aminotransferase; unknown AT, unknown aminotransferase.

### Selection and functional prediction of faba bean TyrA candidates

To investigate the biosynthesis of L-tyrosine in faba bean, specifically the reactions catalyzed by TyrA enzymes, we searched for homologs of the barrel clover TyrA enzymes MtADH (Mt3g071980) and MtPDH1 (AIU94227.1; <u>Schenck *et al*., 2017)</u>. Three candidates showed high homology to both: evgLocus_1024218, evgLocus_220165 and evgLocus_20583 (E-value < 1 x 10^-90^; Data S2). An alignment of the candidates to other characterized legume TyrA proteins shows that all three have characteristic NAD(P)^+^ binding domains (GXGXXG), which are essential for dehydrogenase activity (Fig. 7A; Supplementary Fig. 4). To further assign these candidates, we constructed a phylogenetic tree including known canonical ADHs (chloroplastic, with two dehydrogenase domains), non-canonical ADHs (ncADHs; cytosolic, single domain), and PDHs (cytosolic, single domain; Schenck *et al*. 2015, 2017). The tree showed that evgLocus_1024218 was associated with the canonical TyrA/ADH clade, evgLocus_220165 was associated with ncADH sequences in the non-canonical TyrA clade, and evgLocus_20583 was associated with PDH sequences in the non-canonical TyrA clade (Fig. 7B; Supplementary Fig. 5). To verify this assignment further, we searched for targeting peptides using the online server TargetP-2.0 (Armenteros *et al*., 2019). The results showed that evglocus_1024218 encodes an N-terminal cTP, whereas evgLocus_20583 and evgLocus_220165 do not (Supplementary Table 2). Additionally, we examined the respective residues equivalent to Asp218 of GmncADH, which determines the substrate specificity for arogenate (Fig. 7A; Schenck *et al*. 2017). Both evgLocus_220165 and evgLocus_1024218 (both domains) feature the key Asp residue, whereas evgLocus_20583 has a Cys residue at that position, resembling GmPDH (Fig. 7A). Based on this overall sequence analysis, we predict the following: evglocus_1024218 encodes a canonical, chloroplast-targeted, two-domain ADH hereby dubbed VfADH; evgLocus_220165 encodes a non-canonical, cytosolic ADH hereby named VfncADH; and evgLocus_20583 encodes a cytosolic PDH hereby called VfPDH.

**Figure 7.**
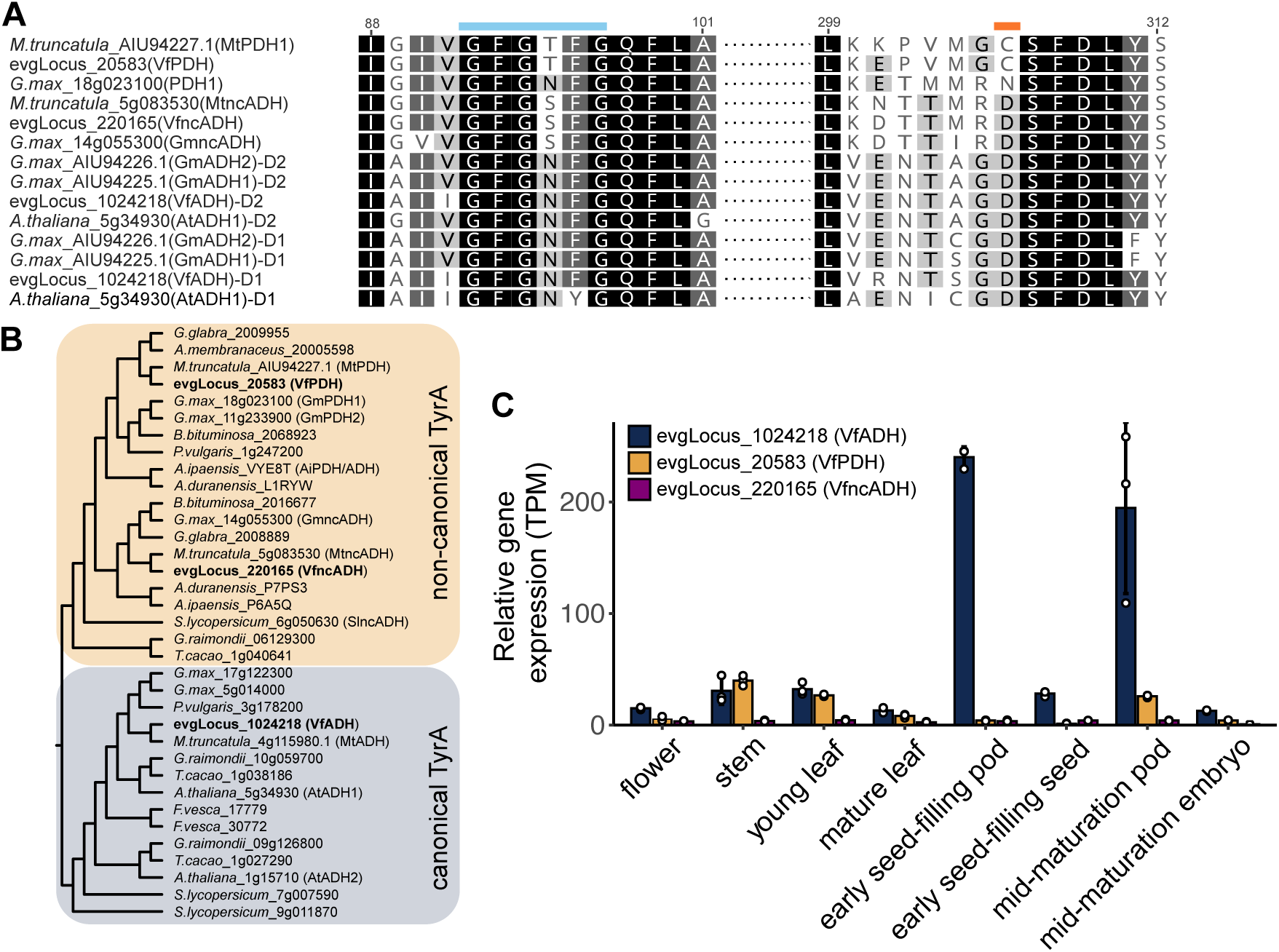
Selection, functional prediction and expression of TyrA candidates in faba bean. (A) Partial protein sequence alignment of the three faba bean TyrA candidates (evgLocus_######) with TyrA enzymes from barrel clover (*Medicago truncatula*), soybean (*Glycine max*) and thale cress (*Arabidopsis thaliana*). The blue line highlights the GXGXXG motif of the NADP+ binding domain. The orange line highlights the active site residue that is crucial for enzyme specificity: Asp for arogenate dehydrogenase (ADH) enzymes; Asn or Cys for prephenate dehydrogenase (PDH) enzymes. The two domains of each ADH were aligned separately (D1, domain 1; D2, domain 2). Numbering is according to the full alignment in Fig. S3. (B) A simplified gene tree (Fig. S4) of TyrA homologs from legume and eudicot lineages, including the three faba bean TyrA candidates (bold). Clade classifications derive from Schenck *et al*. (2017). (C) Expression analysis of VfPDH, VfncADH and VfADH in eight aerial tissues of faba bean. Expression values were obtained from the available transcriptome of the inbred line Hedin/2 (Björnsdotter *et al*., 2021). Hollow circles represent individual data points, bars represent mean values, and error bars represent the standard deviation (±).

To analyze the expression of VfPDH, VfncADH, and VfADH, we turned to the transcriptomic data by Björnsdotter *et al*. (2021), which represents eight aerial tissues of Hedin/2. Inspection of expression patterns showed that VfADH and VfPDH had generally higher expression levels compared to VfncADH, which was lowly expressed in all tissues (Fig. 7C). VfADH was expressed in all tissues, with the highest expression in pods (>100 TPM). By contrast, VfPDH showed highest expression in stems, young leaves, and mid-maturation pods (between 24 and 45 TPM), whereas expression in other tissues was very low (<10 TPM).

### Transient expression of TyrA candidates in *N. benthamiana* in combination with CYP76AD6

We expressed the TyrA candidates transiently in *N. benthamiana*, either on their own or in combination with CYP76AD6 from beetroot (Fig. 8A; Polturak *et al*. 2016). For each combination, we measured L-tyrosine, tyramine (decarboxylation product of L-tyrosine), L-DOPA, and the previously detected L-DOPA hexosides (Fig. 4C). Compared to a GFP control, expression of both VfADH and VfPDH led to a 2-3-fold increase in L-tyrosine (Fig. 8B) and tyramine levels (Fig. 8B), whereas no differences were observed for VfncADH (Fig. 8B). Expression of CYP76AD6 led to accumulation of L-DOPA and its hexosides while at the same time reducing L-tyrosine to undetectable levels (Fig. 8B). Expression of the TyrA enzymes on top of CYP76AD6 did not alter the levels of L-DOPA or dopamine (Fig. 8B), but ADH was able to boost the levels of the L-DOPA hexosides up to 6-fold (Fig. 8B).

**Figure 8.**
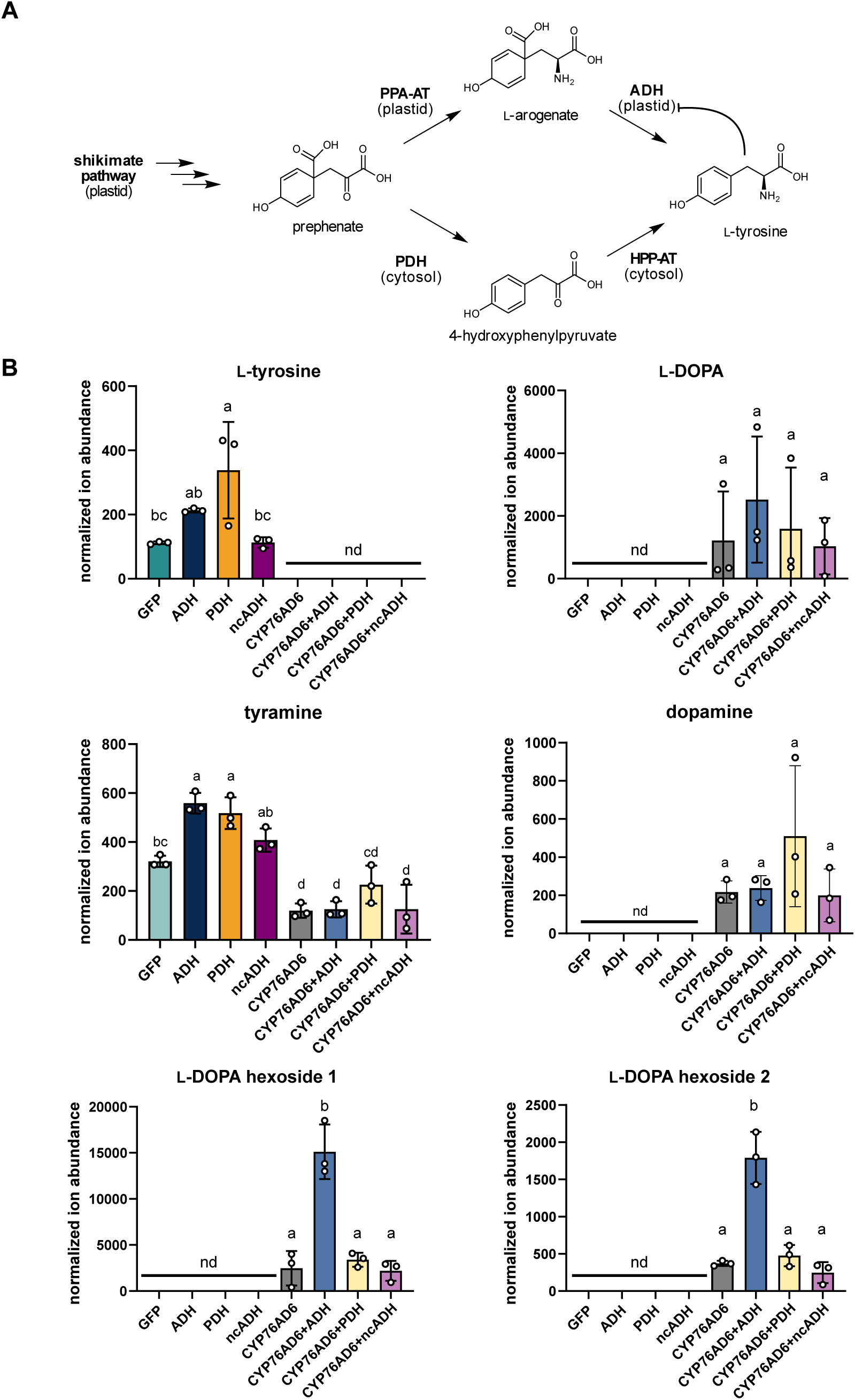
Transient expression of faba bean TyrA genes singly and with beetroot L-tyrosine oxidase, CYP76AD6, in *Nicotiana benthamiana*. (A) The biosynthetic routes to L-tyrosine in legume species via the plastid-localized arogenate dehydrogenase (ADH) or the cytosol-localized prephenate dehydrogenase (PDH). (B) The faba bean TyrA candidates, ADH, PDH and non-canonical ADH (ncADH), were expressed singly or in combination with CYP76AD6 in *N. benthamiana* leaves and metabolites were quantified 5 days post-infiltration. CYP76AD6 was also expressed on its own and GFP was included as a negative control. Each graph shows the quantification of a specified metabolite, normalized to the intensity of the internal standard, caffeine. Hollow circles represent individual data points, bars represent mean values, and error bars represent the standard deviation (±). Letters indicate significant differences in the mean values, identified by one-way ANOVA and Tukey’s post-hoc tests (*p* < 0.05). HPP-AT, 4-hydroxyphenylpyruvate aminotransferase; PPA-AT, prephenate aminotransferase; nd, not detected.

## Discussion

We used different strategies to select candidate genes potentially encoding an L-tyrosine oxidase in faba bean. These included gene-to-metabolite correlation analyses using transcriptomic and metabolomic datasets, as well as homology-based approaches, ultimately testing 15 different CYP, 2-ODD and APX candidate genes. None of the candidate genes led to L-DOPA production when expressed in yeast or *N. benthamiana* (Fig. 4), suggesting that none encoded the oxidase we were searching for.

Correlation-based gene selection strategies rely on the proportional accumulation of metabolites in the tissues where they are synthesized. Accordingly, our correlation-based strategies may have been challenged by possible long-distance transport of L-DOPA across tissues. Transport between tissues is well-known for other L-tyrosine-derived specialized metabolites such as benzylisoquinoline alkaloids in poppy (Dastmalchi *et al*., 2019) and glucosinolates in Brassicaceae species (Du and Halkier, 1998; Chen *et al*., 2001). Importantly, our precursor feeding experiment with L-tyrosine-^13^C9 supports the hypothesis of L-DOPA transport between tissues (Fig. 5). In this experiment, radicles had the highest biosynthetic capacity (Fig. 5B), but it was the aerial tissues that displayed the highest L-DOPA content (Fig. 5C). It is important to note, however, that measuring biosynthetic capacity with feeding experiments can be biased towards tissues that more efficiently absorb and transport the fed precursor into biosynthetic cells. Accordingly, additional experiments more directly showing the hypothesized long-distance transport are needed, for example, grafting experiments with rootstocks and scions displaying different L-DOPA contents (though L-DOPA variation in faba bean accession collections remains unexplored). In any case, our results highlight the possibility of long-distance transport of L-DOPA and suggest using gene discovery strategies different than those based on gene-to-metabolite correlations. Alternative strategies could include carrying out correlation analyses between gene expression and biosynthetic capacity across tissues, testing closer homologues of CYP76AD6 regardless of their expression patterns, or simply focusing on high expression in radicles.

Interestingly, heterologous overexpression of CYP76AD6 in *N. benthamiana* resulted not only in accumulation of L-DOPA but also of dopamine as well as two L-DOPA hexosides (Fig. 4B,C). Dopamine accumulation has been observed before when overexpressing CYP76AD enzymes in *N. benthamiana* (Berman *et al*., 2024; Jung and Maeda, 2024). In this system, dopamine is likely produced by an endogenous tyrosine decarboxylase (TyDC), a conserved enzyme with activity on both tyrosine and L-DOPA (producing tyramine and dopamine, respectively; Torrens-Spence *et al*. 2020). Alternatively, dopamine could be produced through direct oxidation of tyramine by CYP76AD6. To our knowledge, the accumulation of L-DOPA hexosides upon heterologous expression of CYP76AD6 has not been reported previously.

Faba bean seeds accumulate 3′-*O*-β-d-glucopyranosyl-L-DOPA endogenously (Nagasawa *et al*., 1961; Andrews and Pridham, 1965; Irmer *et al*., 2026) whereas maize accumulates an L-DOPA glucoside upon exposure to L-DOPA (Aboshi *et al*., 2023). In maize, glycosylation of L-DOPA is hypothesized to detoxify the molecule (Aboshi *et al*., 2023), as L-DOPA acts as a phytotoxic allelochemical (Matsumoto, 2011; Soares *et al*., 2014). It is likely that endogenous UDP-glycosyltransferases (UGTs) in *N. benthamiana* detoxify L-DOPA in a similar fashion to the ones in maize, given the substrate promiscuity of many UGTs (Aboshi *et al*., 2023; Sirirungruang *et al*., 2025). To date, no naturally occurring L-DOPA glycosyltransferase has been identified.

We also devised and tested a novel hypothesis for the biosynthesis of L-DOPA. The hypothesis was based on the last step of cytosolic L-tyrosine biosynthesis being reversible (Fig. 6A; Maeda and Dudareva 2012; Wang *et al*. 2016), which implies that any L-tyrosine fed to faba bean is likely in equilibrium with its immediate precursor HPP. Under this scenario, HPP could be the direct subject of oxidation (to give 3-OH-HPP), after which a simple transamination would give L-DOPA (Fig. 6A). In this scenario, the transaminase could be either the same one involved in the last step in L-tyrosine biosynthesis, or a homologous, dedicated transaminase. To allow a high accumulation of L-DOPA, this scenario would require L-DOPA to be translocated away from the transaminase, for instance by a dedicated transporter (Fig. 6A). We tested this hypothesis by feeding labeled L-tyrosine (L-tyrosine-^13^C2D5^15^N) to faba bean radicles, with the ^15^N serving as the indicator for whether L-tyrosine would become transaminated en route to L-DOPA (expected loss of ^15^N upon transamination). While there may be some minor amounts of L-DOPA generated via 3-OH-HPP (Fig. 6B), our results show that L-tyrosine does not get transaminated on its way to becoming L-DOPA.

The supply of the precursor, L-tyrosine, is also an important factor affecting L-DOPA accumulation in faba bean. We identified three putative TyrA genes in faba bean: one coding for an ADH (VfADH; evgLocus_1024218), one coding for a non-canonical ADH (VfncADH; evgLocus_220165), and a third coding for a PDH (VfPDH; evgLocus_20583). The identification of a PDH in faba bean corroborates studies showing PDH activity in faba bean tissues (Gamborg and Keeley, 1966; Schenck *et al*., 2020) and supports the cytosolic PDH route for L-tyrosine biosynthesis in faba bean. According to expression in Hedin/2, VfPDH and VfADH have a very similar expression pattern in floral and vegetative tissues (Fig. 7C). However, VfADH has much higher expression levels in fruit tissues, particularly in pods (Fig. 7C). The accumulation of L-DOPA in pods is also very high (Fig. 2B; Fig. S2B), suggesting that VfADH may be key for tyrosine supply in this tissue. On the other hand, L-DOPA accumulation is just as high in flower tissue where VfPDH and VfADH are comparatively low, further suggesting that sites of high L-DOPA accumulation may not correspond with areas of high biosynthetic activity due to potential long-distance transport. Regarding the third TyrA gene, VfncADH, its expression was very low overall thus we hypothesize that it does not contribute to the supply of L-tyrosine for L-DOPA biosynthesis in the analyzed tissues.

Individual expression of the faba bean TyrA genes in *N. benthamiana* revealed that both VfADH and VfPDH could increase the levels of L-tyrosine and tyramine around 2-3-fold (Fig. 8B). Previously, a canonical ADH from beetroot (BvADHβ) was shown unable to mediate such increases, consistent with experimental evidence that it is strongly feedback inhibited by L-tyrosine (Lopez-Nieves *et al*., 2018). In the same publication, a deregulated cognate ADH (BvADHα) was shown to boost L-tyrosine levels 10-fold. It remains to be shown whether our modest increase in L-tyrosine levels as mediated by VfADH corresponds to a relaxation of its feedback regulation properties or to other factors such as the higher levels of recombinant protein expected from our binary vector system (Peyret *et al*., 2019). Interestingly, none of the faba bean TyrA genes were able to increase the levels of L-DOPA or dopamine in *N. benthamiana* upon co-expression with CYP76AD6; however, VfADH boosted the levels of the two L-DOPA hexosides up to 6-fold (Fig. 8B). VfPDH is likely feedback insensitive, like other characterized PDHs (Schenck *et al*., 2015, 2017), but we cannot attribute the observed increase to this feature. Instead, the marked difference between VfADH and VfPDH may come down to sub-cellular localization and the ability of different sub-cellular compartments in *N. benthamiana* to support a high flux towards L-tyrosine. While VfADH is predicted to be plastidic, VfPDH lacks a targeting peptide and is likely cytosolic (Table S2). Cytosolic L-tyrosine biosynthesis via PDH (prephenate dehydrogenase) relies on a cytosolic source of prephenate as well as on the availability of a cytosolic aminotransferase (Schenck *et al*., 2015, 2017). While some elements of a cytosolic aromatic amino acid pathway likely exist in *N. benthamiana* (Sommer and Heide, 1998; Qian *et al*., 2019), the extent to which it can support a high flux remains unknown. From our experiments, it is clear that VfADH can enable such high flux via plastidic L-tyrosine biosynthesis.

Note that our experiments in *N. benthamiana* do not provide information on whether the PDH-dependent cytosolic route or the ADH-dependent plastidic route are responsible for supporting L-DOPA biosynthesis in faba bean. To investigate this, functional genetic studies in faba bean are needed. Analysis of a barrel clover PDH mutant, for instance, has shown that the cytosolic PDH route is not the principal route for L-tyrosine biosynthesis, but its contribution becomes evident upon feeding with shikimate precursor (metabolic push) (Schenck *et al*., 2020). In *Inga* species, high PDH expression is correlated to hyper-accumulation of L-tyrosine-derived specialized metabolites such as Tyr-gallates (Coley *et al*., 2019). Elucidation of the L-tyrosine oxidase enzyme in faba bean and its testing via functional genetics together with the TyrA genes here identified will help determine if the cytosolic L-tyrosine route is important not only upon high metabolic push, but also under conditions of high metabolic pull.

## Conclusions

Our study narrows down the pool of candidates for the L-tyrosine oxidase in faba bean and highlights the possibility of long-distance transport of L-DOPA, which should be considered in future candidate selection strategies. Precursor feeding studies unveiled the high biosynthetic activity of faba bean radicles and disproved an alternative biosynthetic hypothesis via the pathway intermediate HPP. Our study also uncovered three putative TyrA genes in faba bean. From these, VfADH and VfPDH can increase the levels of L-tyrosine 2-3-fold when expressed in *N. benthamiana*. Finally, VfADH can uniquely boost the levels of L-DOPA-derived hexosides up to 6-fold in the same system. Our work contributes significantly to the search for L-tyrosine oxidase enzymes and kickstarts the characterization of L-tyrosine and L-DOPA biosynthesis in faba bean.

## Acknowledgements

We thank Adam Jozwiak and Asaph Aharoni (Weizmann Institute of Science, Israel) for providing a clone of CYP76AD6, Hadrien Peyret and George Lomonossoff (John Innes Centre, United Kingdom) for providing empty pHREAC and pHREAC-GFP vectors, and John Dueber (University of California, Berkeley, USA) for the yeast strain yWCD230.

## Author Contributions

X.X. and F.G.-F. conceived the research plan and developed the project design. X.X., H.St., D.B., D.M., B.D. L.E-H, and H.Sh. carried out the experiments and data analysis. F.G.-F., S.A. and H.Sh. provided the resources and acquired the funding. F.G.-F coordinated the project. X.X., H.St., B.D, and H.Sh. prepared the figures. X.X., H.St., D.B., F.G.-F. and H.Sh. wrote the manuscript with input from all authors.

