## Supplementary information for "Exploring L-tyrosine and L-DOPA biosynthesis in faba bean (*Vicia faba* L.)"

### Authors contributed equally

*Authors for correspondence:

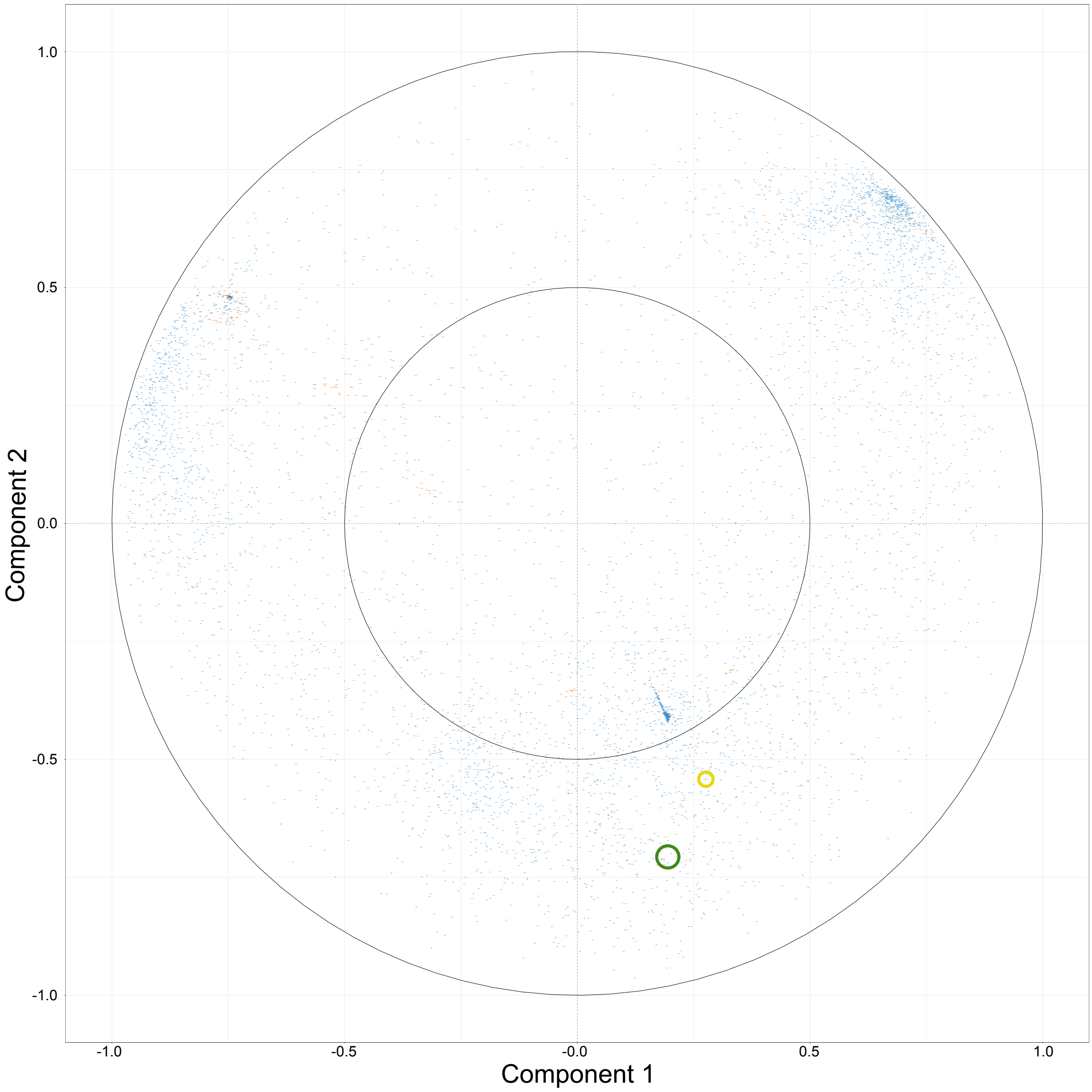


**Figure S1**. A correlation circle plot combining the gene expression dataset and the metabolomics dataset from eight aerial tissues of faba bean using the partial least squares regression method. Physical closeness indicates a high degree of correlation. Genes are represented as blue dots and metabolic features as orange dots. The metabolite features associated with L-DOPA are circled in dark green and the gene candidate closely correlated with these features, evgLocus_256923, is circled in yellow.

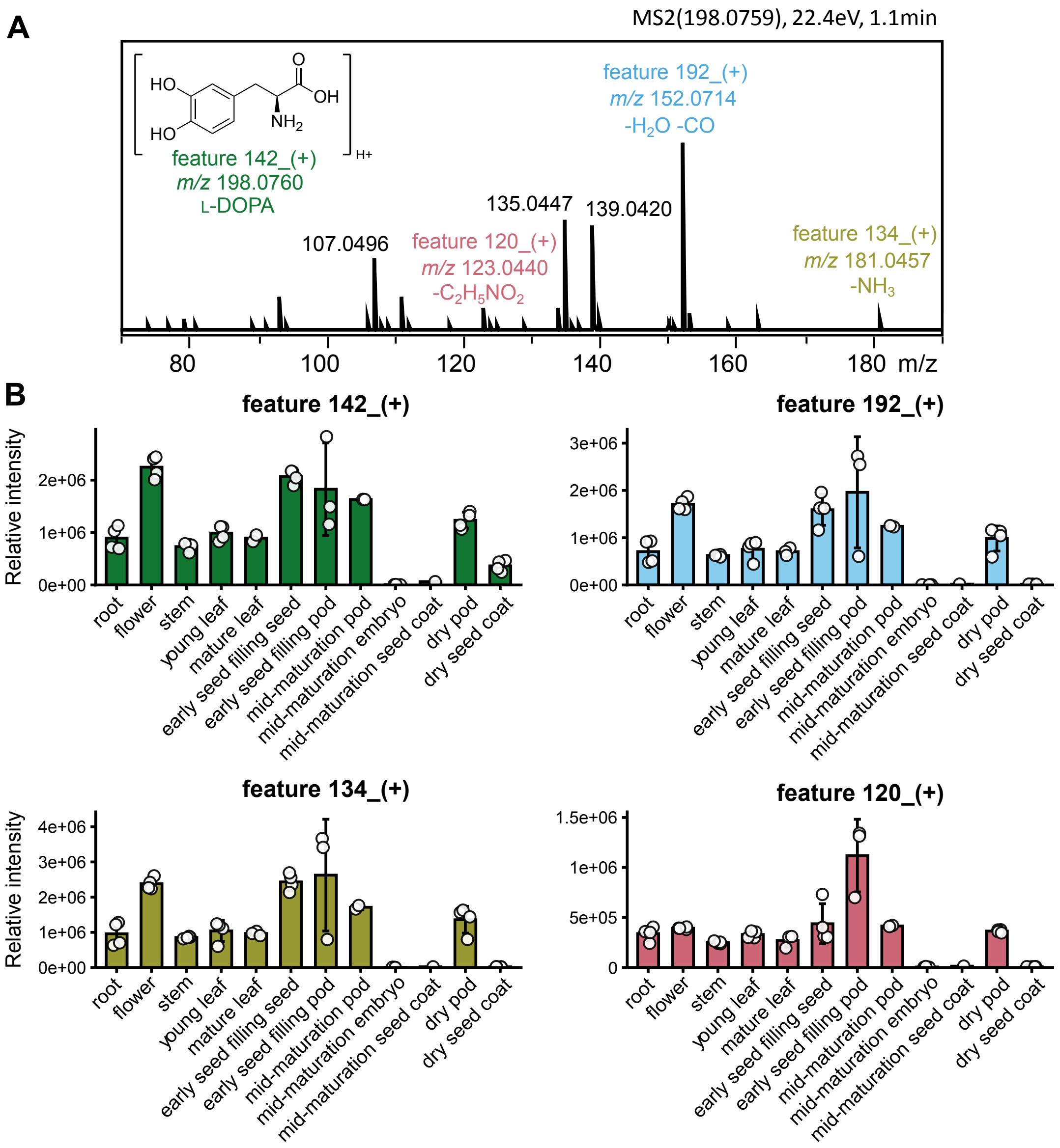


**Figure S2.** Metabolic features representing L-DOPA and their abundance in twelve faba bean tissues as inferred from the expanded metabolomic dataset. (**A**) MS2 spectra of a commercial standard of L-DOPA. The selected metabolic features are indicated above the corresponding daughter ion signals (colored labels). (**B**) Relative levels of L-DOPA in different tissues of faba bean as indicated by the mean abundance of L-DOPA-associated metabolic features across faba bean tissues. The intensities of metabolic features were normalized to the internal standard (caffeine) and to the dry weight of each sample. Hollow circles represent individual data points, bars represent mean values, and error bars represent the standard deviation (±).

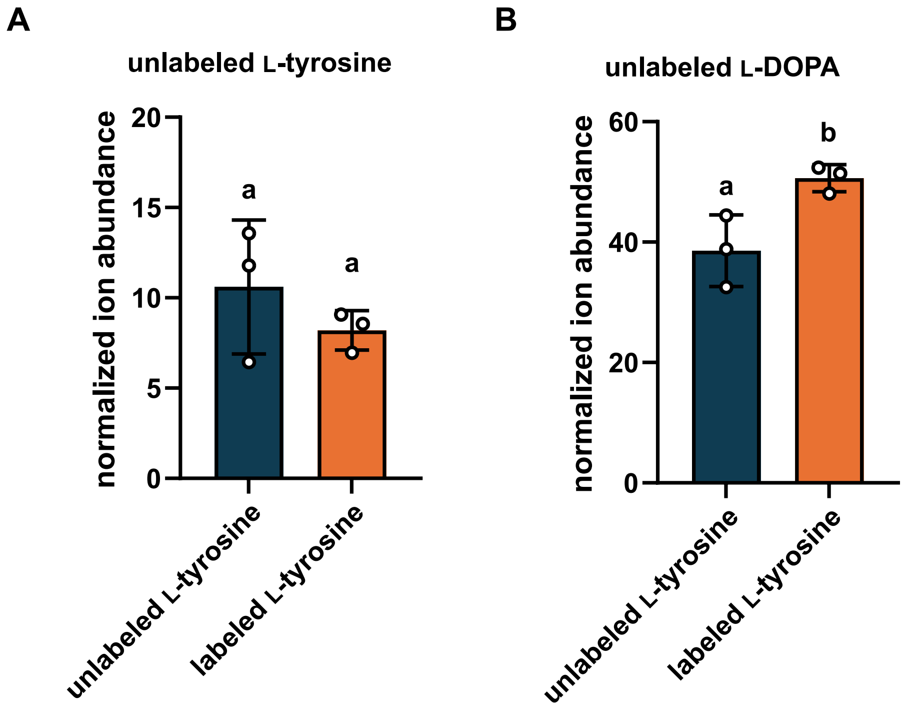


**Figure S3**. Quantification of unlabeled L-tyrosine and unlabeled L-DOPA in feeding experiments in seedling tissues of faba bean. (B) LC-MS quantification of unlabeled L-tyrosine and unlabeled L-DOPA from radicles, seedling stems, seedling leaves, and seedling roots that were fed with either unlabeled L-tyrosine (blue bars) or L-tyrosine-$^{13}C_2D_5{}^{15}N$ (orange bars). Quantification was carried out relative to the internal standard, caffeine. Hollow circles represent individual data points, bars represent mean values, and error bars represent the standard deviation ($\pm$). Letters indicate significant differences in the mean values, identified by two-way ANOVA and Tukey's post-hoc tests ($p < 0.05$).

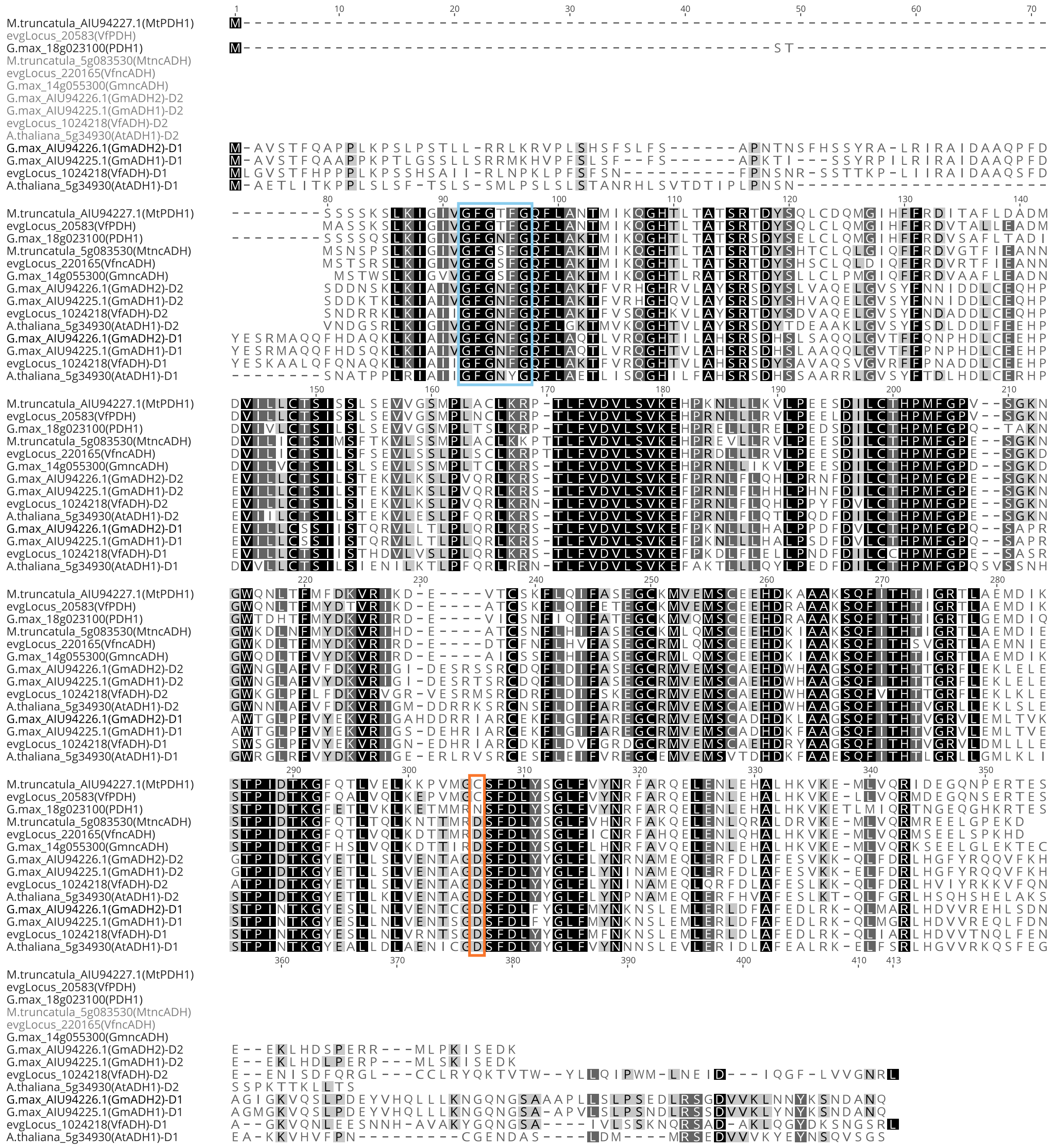


**Figure S4**. Protein sequence alignment of candidate faba bean TyrA sequences together with characterized TyrA sequences from other legumes. The blue box highlights the GXGXXG motif of the NAD(P)$^+$ binding domain. The orange box indicates the active site residue that is crucial for enzyme specificity: Asp for arogenate dehydrogenase (ADH) enzymes, and Asn or Cys for prephenate dehydrogenase (PDH) enzymes. The reference sequences are GmADH1 from soybean (Glyma5g07870), GmADH2 from soybean (Glyma17g13150), AtADH1 from thale cress (NP_198343.1), MtPDH from barrel clover (3g071980), GmPDH1 from soybean (18g02650), MtncADH from barrel clover (5g083530), and GmncADH from soybean (14g05990). The two domains of each canonical ADH and the putative faba bean ADH were aligned separately (D1, domain 1; D2, domain 2).

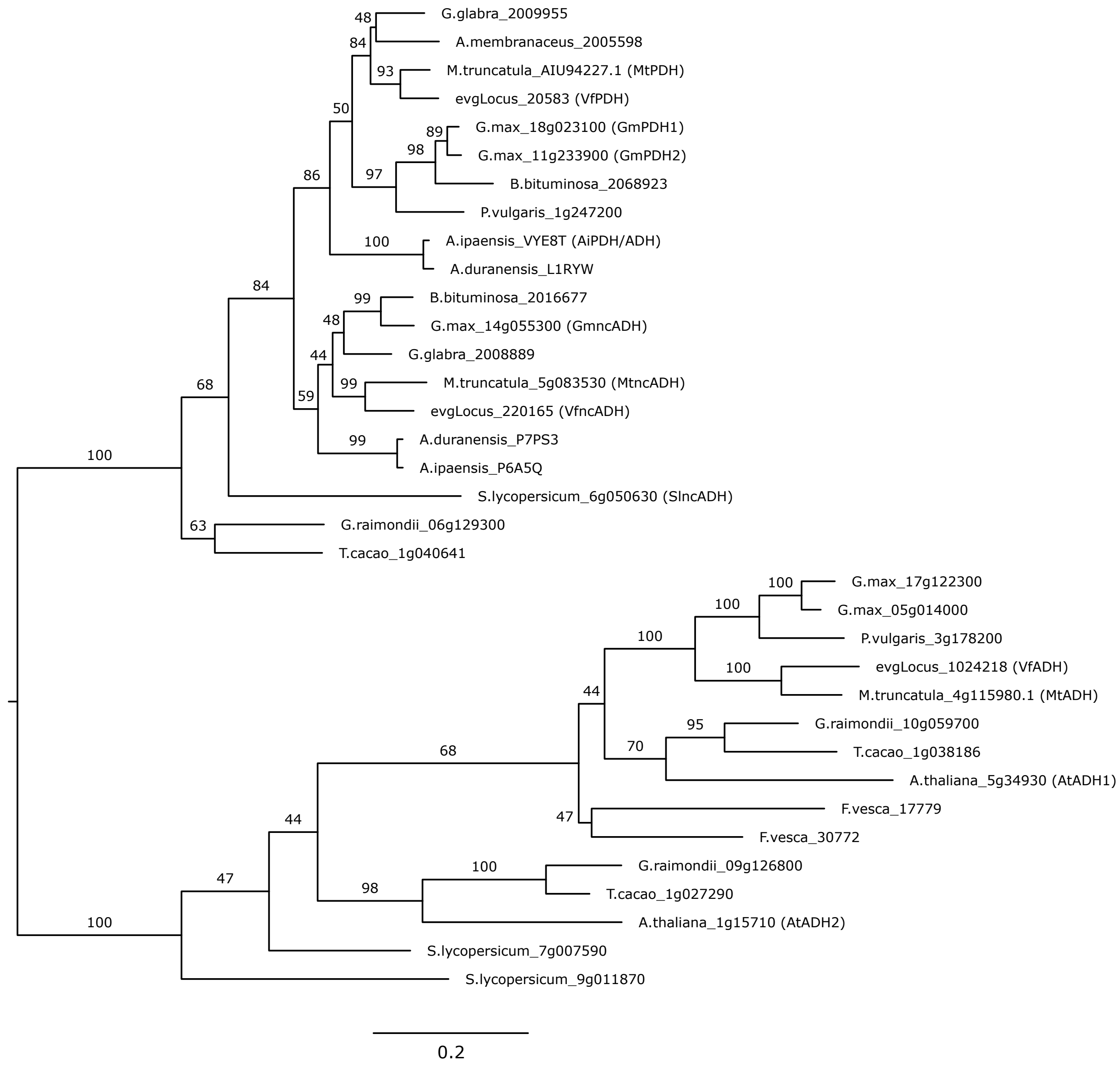


**Figure S5**. TyrA gene tree with amino acid branch lengths and support values. A maximum liklihood tree was inferred from an amino acid sequence alignment of the faba bean candidate TyrA enzymes and TyrA sequences from Fig. 1b of Schenck *et al*. (2017) with the addition of arogenate dehydrogenase from barrel clover (MtADH; Mt3g071980). The Whelan and Goldman model was used to estimate sequence evolution and branch lengths are expected substitutions per site. Branch values show support from 1000 non-parametric bootstraps.

**Table S1.** Oligonucleotides used in this study.

| Name | Sequence (5' to 3') | Purpose |
|---|---|---|
| HS-TUW105 | CACCACAGGTCTCGTATGATACA ACTATTCCTCTTTTCACACTCA | Cloning evgLocus_40272 into pEVY002 |
| HS-TUW106 | CACCACAGGTCTCGGGATCCTGT TGCTTCCCCGTGAA | Cloning evgLocus_40272 into pEVY002 |
| HS-TUW107 | CACCACAGGTCTCGTATGTGTTC AAGTGCGAGTAACCGT | Cloning evgLocus_1209062 into pEVY002-part1-F |
| HS-TUW108 | CACCACAGGTCTCGCCTCACGG AGCGAAAGTTTAC | Cloning evgLocus_1209062 into pEVY002-part1-R |
| HS-TUW109 | CACCACAGGTCTCGGAGGCCCA ACGACCATGTTTCTTCC | Cloning evgLocus_1209062 into pEVY002-part2-F |
| HS-TUW110 | CACCACAGGTCTCGGGATCCAA CACTTTGGTACACAAA | Cloning evgLocus_1209062 into pEVY002-part2-R |
| HS-TUW111 | CACCACAGGTCTCGTATGGCATT TTCAACACATGAGCAACA | Cloning evgLocus_1273529 into pEVY002 |
| HS-TUW112 | CACCACAGGTCTCGGGATCCTTG AGTCACACGGTACCT | Cloning evgLocus_1273529 into pEVY002 |
| HS-TUW113 | CACCACAGGTCTCGTATGCTAGC GATAATCTGTTCCAGT | Cloning evgLocus_1282817 into pEVY002 |
| HS-TUW114 | CACCACAGGTCTCGGGATCCGA AAGTGCTGGCTGCATA | Cloning evgLocus_1282817 into pEVY002 |
| HS-TUW117 | CACCACAGGTCTCGTATGGATTA TGCGATTAGTGCACTGCT | Cloning evgLocus_366734 into pEVY002 |
| HS-TUW118 | CACCACAGGTCTCGGGATCCTAA CACTTTGTTTAGTGCAACC | Cloning evgLocus_366734 into pEVY002 |
| HS-TUW119 | CACCACAGGTCTCGTATGACTCT GTTTCTCACCATACCTCT | Cloning evgLocus_727341 into pEVY002- part1-F |
| HS-TUW120 | CACCACAGGTCTCGGGATCCGAT ATCAGCTGGCACATG | Cloning evgLocus_727341 into pEVY002-part3-R |
| HS-TUW123 | CACCACAGGTCTCGTATGAGAGA ACTTGTGCACAAATATGA | Cloning evgLocus_256923 into pEVY002 |
| HS-TUW124 | CACCACAGGTCTCGGGATCCGTT ACTAGTAGGAGTTGCA | Cloning evgLocus_256923 into pEVY002 |
| HS-TUW125 | CACCACAGGTCTCGACGGCGAG CACCGTG | Cloning evgLocus_727341 into pEVY002-part1-R |
| HS-TUW126 | CACCACAGGTCTCGCCGTGACC GACTTACTCGAGCCAT | Cloning evgLocus_727341 into pEVY002-part2-F |
| HS-TUW127 | CACCACAGGTCTCGTCCAATGAT GGTGTCTTCACT | Cloning evgLocus_727341 into pEVY002-part2-R |
| HS-TUW128 | CACCACAGGTCTCGTGGACTCCT TTGGGACATGATAACTGC | Cloning evgLocus_727341 into pEVY002-part3-F |
| HS-TUW129 | CACCACAGGTCTCGTATGGGAAA ATCTTACCCTACCGT | Cloning evgLocus_959121 into pEVY002 |
| HS-TUW130 | CACCACAGGTCTCGGGATCCGG CTTCGGCAAATCCAAG | Cloning evgLocus_959121 into pEVY002 |
| HS-TUW131 | CACCACAGGTCTCGTATGGTAGA ACGTGTGTTACTCACTCCT | Cloning evgLocus_89125 into pEVY002 |
| HS-TUW132 | CACCACAGGTCTCGGGATCCGTT TCCAAGCAGGGATGT | Cloning evgLocus_89125 into pEVY002 |
| HS-TUW133 | CACCACAGGTCTCGTATGGCGTT GCCGGTGGTGG | Cloning evgLocus_145931 into pEVY002 |
| HS-TUW134 | CACCACAGGTCTCGGGATCCCTT CCCTCTTCTGGAATGT | Cloning evgLocus_145931 into pEVY002 |
| HS-TUW135 | CACCACAGGTCTCGTATGGCTCA ATATAGAGTTGACGCAGA | Cloning evgLocus_255817 into pEVY002 |
| HS-TUW136 | CACCACAGGTCTCGGGATCCATA ATTCCCGAAAATAAGAGT | Cloning evgLocus_255817 into pEVY002 |
| HS-TUW137 | CACCACAGGTCTCGTATGACATT GCCATTATCATCCATGGA | Cloning evgLocus_1242091 into pEVY002 |
| HS-TUW138 | CACCACAGGTCTCGGGATCCATC CCTGTGTAATGCTTTGA | Cloning evgLocus_1242091 into pEVY002 |
| HS-TUW139 | CACCACAGGTCTCGTATGGAAAT TTTTGAGATAAGCACA | Cloning evgLocus_30215 into pEVY002 |
| HS-TUW140 | CACCACAGGTCTCGGGATCCGA CTTTTGTCATGTTGAGAACA | Cloning evgLocus_30215 into pEVY002 |

| Name | Sequence (5' to 3') | Purpose |
|---|---|---|
| HS-TUW143 | ACACTATATCAATAAAGATCTATG GCACAATATAATAGAGTTGATGC TGAGT | HiFi cloning evgLocus_145163 into pEVY002 |
| HS-TUW144 | TTCGCCTCGAGTTAGGATCCGTT GATTTTTTTGTTGAGTTCA | HiFi cloning evgLocus_145163 into pEVY002 |
| HS-TUW145 | ATCCTAACTCGAGGCGAATTTC | HiFi cloning into pEVY002 plasmid backbone |
| HS-TUW146 | CATAGATCTTTATTGATATAGTG | HiFi cloning into pEVY002 plasmid backbone |
| evgLocus_1242091_F | GGCTTAA[U]ATGACATTGCCATT ATCA | Cloning into pEAQ-USER |
| evgLocus_1242091_R | GGTTTAA[U]CTAATCCCTGTGTA ATGCT | Cloning into pEAQ-USER |
| evgLocus_30215_F | GGCTTAA[U]ATGGAAATTTTGA GATAAGCACA | Cloning into pEAQ-USER |
| evgLocus_30215_R | GGTTTAA[U]TCAGACTTTTGTCA TGTTGAGAAC | Cloning into pEAQ-USER |
| evgLocus_366734_F | GGCTTAA[U]ATGGATTATGCGAT TAGTGC | Cloning into pEAQ-USER |
| evgLocus_366734_R | GGTTTAA[U]TTATAACACTTTGTT TAGTGCAAC | Cloning into pEAQ-USER |
| evgLocus_1282817_F | GGCTTAA[U]ATGCTAGCGATAAT CTGTTCCA | Cloning into pEAQ-USER |
| evgLocus_1282817_R | GGTTTAA[U]TTAGAAAGTGCTGG CTGCATA | Cloning into pEAQ-USER |
| evgLocus_1273529_F | GGCTTAA[U]ATGGCATTTTCAAC ACATG | Cloning into pEAQ-USER |
| evgLocus_1273529_R | GGTTTAA[U]CTATTGAGTCACAC GGTACCTAT | Cloning into pEAQ-USER |
| evgLocus_256923_F | GGCTTAA[U]ATGAGAGAACTTGT GCACAAATAT | Cloning into pEAQ-USER |
| evgLocus_256923_R | GGTTTAA[U]TTAGTTACTAGTAG GAGTTGCA | Cloning into pEAQ-USER |
| evgLocus_40272_F | GGCTTAA[U]ATGATACAACTATT CCTCTTTTTC | Cloning into pEAQ-USER |
| evgLocus_40272_R | GGTTTAA[U]TCATGTTGCTTCCC CGTGAA | Cloning into pEAQ-USER |
| evgLocus_959121_F | GGCTTAA[U]ATGGGAAAATCTTA CCCTACCG | Cloning into pEAQ-USER |
| evgLocus_959121_R | GGTTTAA[U]TTAGGCTTCGGCAA ATCCA | Cloning into pEAQ-USER |
| evgLocus_89125_F | GGCTTAA[U]ATGGTAGAACGTGT GTTACTCA | Cloning into pEAQ-USER |
| evgLocus_89125_R | GGTTTAA[U]TTAGTTTCCAAGCA GGGATGTC | Cloning into pEAQ-USER |
| evgLocus_145163_F | GGCTTAA[U]ATGGCACAATATAA TAGAGTTGAT | Cloning into pEAQ-USER |
| evgLocus_145163_R | GGTTTAA[U]TCAGTTGATTTTTTT GTTGAG | Cloning into pEAQ-USER |
| evgLocus_1209062_F | GGCTTAA[U]ATGTGTTCAAGTGC GAGTAA | Cloning into pEAQ-USER |
| evgLocus_1209062_R | GGTTTAA[U]TTAAACACTTTGGT ACACAAATT | Cloning into pEAQ-USER |
| evgLocus_145931_F | GGCTTAA[U]ATGGCGTTGCCGGT GGTGGTTG | Cloning into pEAQ-USER |
| evgLocus_145931_R | GGTTTAA[U]CTACTTCCCTCTTCT GGAATGTTC | Cloning into pEAQ-USER |
| evgLocus_255817_F | GGCTTAA[U]ATGGCTCAATATAG AGTTGAC | Cloning into pEAQ-USER |
| evgLocus_255817_R | GGTTTAA[U]TCAATAATTCCCGA AAATA | Cloning into pEAQ-USER |

| Name | Sequence (5' to 3') | Purpose |
|---|---|---|
| evgLocus_297956_F | GGCTTAA[U]ATGACCTCAAACAA ACCACA | Cloning into pEAQ-USER |
| evgLocus_297956_R | GGTTTAA[U]TTACAACTTGAGTT TAGTAATGTCG | Cloning into pEAQ-USER |
| evgLocus_727341_F | GGCTTAA[U]ATGACTCTGTTTCT CACCATAC | Cloning into pEAQ-USER |
| evgLocus_727341_R | GGTTTAA[U]TTAGATATCAGCTG GCACATGT | Cloning into pEAQ-USER |
| ncADH_pHREAC_GA_F | ctacattatattaaacgtctctATGTCTACTTC ACGGTCTCTGA | Cloning evgLocus_220165 into pHREAC |
| ncADH_pHREAC_GA_R | tttaatgaaaccagagcgTCAGTCATGTTT GGGACTCAGC | Cloning evgLocus_220165 into pHREAC |
| ADH_pHREAC_GA_F | ctacattatattaaacgtctctATGCTCGGCG TTTCAACCTT | Cloning evgLocus_1024218 into pHREAC |
| ADH_pHREAC_GA_R | tttaatgaaaccagagcgTCATTTAGCATC CACGGTATCTGT | Cloning evgLocus_1024218 into pHREAC |
| PDH_pHREAC_GG_F | caccacaggtctcgaaaaATGGCATCTTC CAAAAGTTTGA | Cloning evgLocus_20583 into pHREAC |
| PDH_pHREAC_GG_R | caccacaggtctcgagcgTCAACTTTGAG TTCTTTCTGAGT | Cloning evgLocus_20583 into pHREAC |
| CYP76AD6_pHREAC_F | CACCACAGGTCTCGAAAAATGG ATAACGCAACACTTGCTG | Cloning CYP76AD6 into pHREAC |
| CYP76AD6_pHREAC_R | CACCACAGGTCTCGAGCGCTAG TTTCTGGGAACTGGAA | Cloning CYP76AD6 into pHREAC |

**Table S2**. Probability of sub-cellular localization of faba bean TyrA enzymes (VfADH, VfncADH and VfPDH), inferred using TargetP-2.0.

| TyrA candidate | Prediction | OTHER | SP | mTP | cTP | lTP |
|---|---|---|---|---|---|---|
| evgLocus_20583 (VfPDH) | OTHER | 0.937 | 0.027 | 0.013 | 0.021 | 0.002 |
| evgLocus_220165 (VfncADH) | OTHER | 0.870 | 0.022 | 0.099 | 0.006 | 0.003 |
| evgLocus_1024218 (VfADH) | cTP | 0.021 | 0.000 | 0.000 | 0.973 | 0.006 |

SP, signal peptide; mTP, mitochondrial target peptide; cTP, chloroplast transit peptide; lTP, thylakoidal lumen composite transit peptide. Localizations were predicted using TargetP-2.0 (https://services.healthtech.dtu.dk/services/DeepLoc-2.1/).

**Appendix S1.** Coding sequences of gene candidates for L-tyrosine oxidase

>evgLocus_1242091
ATGACATTGCCATTATCATCCATGGATTTTCTATCAAATCCTTATCTTTTTGGAGCTTTGTTTGCATCTTTAACTCTCTT
AATACTTCAATTTCTATTAAAAAACAATAAGAAATATCATCCTGTTGCTGGAACTGTATTCAATCAGTTGATGAACTTCA
ACACACTTCATCATTACATGACTGATCTTGCAAGGAAATACAAGACATACAGGATTCTTAACCTTTTCAGAAATGAAGTT
TATACTGCAGAACCAAACAATGTTGAGTATATACTCAAAACCAACTTTGACAATTATGGAAAGGGGTTGTACAACTATCT
CAATTTGAAGGATTTACTCGGAGATGGGATTTTCACGGTCGATGGCGAGAAATGGCGCGAACAAAGGAAGATATCAAGTC
ATGAATTCTCCACCAAGATGTTAAGGGATTTCAGTACTTTAATATTCAGAAAGAATGCAGCAAAAGTTGCAAATATTGTG
TCTGAAGCTGCAATTTCTAATACTAAGTTAGAAATTCAAGATCTTTTCATGAAATCAACACTGGATTCAATTTTCAAAGT
TGCGTTTGGAACTGAACTTGACAGCATGTGTGGAACAAATGAAGAAGGGAAGAATTTCGCCAATGCATTTGATTCTGCAA
GTGCGTCAACGCTTTATCGTTATGTTGATGTCTTTTGGAAGATAAAGAAGTTTCTAAATATTGGATCAGAGGCAGAATTA
AAGAAAAACACTCAAATCTTAAATGAATTTGTCATTAAGCTAATCAACATCAGAATTCACCAAATGAAGAATTCAAAGGG
TGATTCTATTAGAAAAAGTGGAGATATTCTCTCGAGGTTTCTGCAAGTGAAGGAATATGATACAACATACTTGAGAGATA
TAATTATAAACTTTGTTATTGCGGGGAAAGACACGACGGCCGCTACGCTTTCTTGGTTCGTTTACATGCTATGCAAGTAT
CCTACAGTACAAGAAAAGGTTGCAGAAGAAGTGAGAGAAGCGACAAACACAAAAACAATTAGTAGCTACACTGAGTTTGT
TTCAAGGGTAACAGATGAAGCTATTGAAAAGATGAATTATCTCTATGCAGCACTCACAGAAACTCTCAGACTTTATCCTG
CAGTTCCTGTGGATGCAAAGATTTGTTTTTCTGATGATACATTACCAGATGGATATAGTGTAAAAAAAGGAGACATGGTG
TGTTACCAACCTTATGCAATGGGGAGGATGAAATTCATATGGGGTGATGATGCCGAGGAGTTTAGACCAGAAAGATGGCT
TGATGAGAATGGAGTTTTTCAGCCAGAAAGCCCTTTCAAGTTCACTGCTTTTCAGGCAGGTCCTCGGATATGCCTAGGAA
AAGAGTTTGCTTATAGACAGATGAAGATATTCTCAGCTGTTTTATTAGGTTGTTTTCGTTTCAAATTGAATGATGAGAAA
AAAAATGTGACTTATAAGACAATGATAAATTTTCATATTGATGGAGGTCTTGAAATCAAAGCATTACACAGGGATTAG
>evgLocus_30215
ATGGAAATTTTTGAGATAAGCACAATAATGAAAGTGTTGTTTGGAGCAGTTTTGGTGATTTTGGTTCATATTTTGAATAT
TTTGGTGTTGAGTCCAAAATCACTTAGAGCAAAGCTTCATAGACAGGGTATTAATGGTCCTTCTCCTCACTTCTACTTTG
GCAATATTCCAGAAATAAAGAGACTTTTACTCCAAACTCAATCATCACAAGAAAACAACAAAGATTGTGTTTCAACATCC
ATTTCTCATGACTGGCATTCTAATCTCTTCCCTCATATTTACAAGTGGAAAAAACAATACGGTCCCATTTTCTTGTTTTC
TTCTGGGAGCATACAATGGTTACTGGTGACAGATATAGATATGGTGAAGGAAATTGTCCTACACACGTCTTTGAATCTTG
GGAAGCCTTCTTTTATGTCCAGAGATAATAGGCCACTCCTTGGACAAGGTATATTATCATCAAGTGGTCAATATTGGGCT
CATCAAAGGAAGATAATTGCCCCTGAACTGTACCTAGAGAAAATGAAGGCAAAGGTCAATATGATAATGGACTCAACAAA
TGGACTACTAAGATTATGGGATACCAAATTGAAAAACCATGGAGACAGTTTAGAGATTAAAATTGATCAAGACTTGCGAA
ACTTATCATCTGACATAATTGCAAAAGCTTGTTTTGGAAGCAATTATGTTGAAGGAAGAGAAATATTCACAAAGCTTAGA
GAACTTCAAAACGTCATTTCCAAGATATTTGCTGGAATTCCAGGCTATAGATATTTGCCAAACAAAACCAATAGACAAAT
GTGGAAATTAGAAAAAGAGATAAACTCCAAGATATCAAACCTCATAAAGAAACGCCAAAACAATGCTCGAGATGAACAAG
ATCTCTTGCAAATGATACTCGACAGCGCAAAGAAATGCAAAAGTGGTGATAGTTTCTTGCCGAATTCGATCTCTCAAGAC
ATATTTATAATCGATAATTGCAAAAATATTTTTTTTGCTGGATATGAGACTACTGCAATTACAGCATCATGGTGCTTAAT
GTTGTTATCTACACACCCTGATTGGCAAGATCGTGTTCGCGCCGAGGTGCTTGAAGTTGTTGGAAAAGATGGTAATATAG
ATGCAACTATGCTTAAAAGCTTGAAAATGTTAACTATGGTGATTCAAGAGACATTGAGGCTTTATCCACCAGCGTCATCG
GTTAATCGAGAAGTTTTCAAAGACATTGTTTTCAAAGGCCTATTAGTTCCGAAAGGGATGAATATTCACATTCCAATGCC
GATTTTGCATCAAGACTCGAAGCTTTGGGGGGATGATGCACATGAATTTAATCCGGAAAGATTTGCAAATGGAGTGCAAG
GAGCATGCAAGATTCCACAAGTTTACATGCCATTTGGAATGGGGTCTCGTGTATGTGTAGGACAGCATTTGGCCATGGTT
GAGTTGAAGGTGATGTTGTCTCTCATTCTGTTGGAGTTTCGGTTTTCAGTGTCGCCGAGTTATCGTCATTCGCCTTCCTT
CCACATGCTTATTGAGCCCGGTCATGGAGTTGTTCTCAACATGACAAAAGTCTGA
>evgLocus_40272
ATGATACAACTATTCCTCTTTTTCACACTCACCTTCATTCTCATCATCACCACCACCTTCATCCTCCGTTCAAAACAGCC
CAAGCATGACAGAAAACACCCACCAGGTCCACCAGGCTATCCTCTCATTGGTAACCTCCACATGTTAGGCACCCTCCCAC
ACCGTTCCCTCGAAGCCTTATCCAAAAAACATGGTCCCATCATGTCACTCCGTTTAGGCCAAGTCCCAACAGTCATAGTC
TCTTCCTCTTCAGCAGCCGAACAATTCCTCAAAGCAAACGACCTCGCTTTCGCCAGCCGTCCTAAACTCGAAGCCACGCA
CTACCTATCTTACGGTTCTAAGGGGTTAATTTTTGCTGAATATGGCGCGTATTGGCGTAACATGAGGAAGATTTGCACTC
TGCAGCTTTTAAGTGCTTCTAAAGTTGAGACGTTTGCGTCTTTGAGGAAGAGAGAAATGGAGTTAGCTTTGAAGTTGTTG
AAGAAAGCTGCGAGTTCGGGTGAGGTTGTTGATGTGAGTGAGTTTGTGCATGATGTTATAATGGATATTGTGTGTAAGAT
GGTGTTGGGGTGTAGTATTGATGAGGTTTTTGATTTGAAGAGGCTGATTCAACAAGGGACGAACCTTTCTGGTGCTTTTA
ATCTTGCTGATTATGTTCCTTTTCTTCGAGTTTTTGATCTTCAGGGTATAAGAAGAAGATACAAGAGAACTCATAAAGAA
CTTGACCAAGTCTTGGAGAAGATAGTTAAGGAGCATAAAGAAAGTTCAGATGTACAGAATGGAGGACAGAAGCATAAGGA
CTTTATAGACATTCTACTGTCACAGATGCACCAGCCGATCGGTCCCTCTGATGAGCAAAACAACGTCGTTGATCGAACAA
ATATAAAGGCTATAGTCTTAGACATGATTGCTGCTGCATTCGAAACTTCTGCTACCGTCGTGGAGTGGGCTTTATCCGAG
CTCATGAGACATCCAAGAGTAATGAAAAATCTCCAACAAGAGTTAGATAATGTGGTTGGGGTCAACAAAGTGGTGGAGGA
AAATGATTTGTCAAAGTTAAGTTACTTAGATATTGTGATCATGGAGACGTTAAGATTATATCCCGCAGGGCCACTCGTGC
CACGCGAGTCAACGGAGGATGCTACAGTTGATGGTTATTTCCTAGAAAAGAAATCAAGAATCATTGTGAATTTATGGGCG
ATAGGAAGAGATTCTAAGATATGGTCCGACAATGCGGAAGAATTTTATCCAGAGAGATTCATGGATAAAAATTTAGATTA
CCGCGGAAATGATTTCCAGTTTATACCATTTGGTTTTGGTCGAAGAGGATGTCCTGGAATAAACATGGGACTAATTACAG
TGAAACTTGTTGTCACTCAACTGGTACATTGTTTTTCTTGGAAACTTCCCTCGAATATGACTCCAAATGATTTGGACATG
ACAGAAAAAATTTGGTCTTTCAATCCCAAGAGCCAAACACTTGCTTGCGGTGCCAAAATATCGTCTTCACGGGGAAGCAAC
ATGA
>evgLocus_89125
ATGGTAGAACGTGTGTTACTCACTCCTTCACTCCCATTACCATCTCGACCTTCACTTTCACCAACCACAATGGCTACTAT
TACTGGCGGCGCCGCCACTCGTATCATCCCCTCCGCCACCAGAGCCACCGTCTCACTCTCCTCTTCTTCCCGTTCCTTTT
TCTCATTTTCATCCTCTTCTACTTCCGTTTCCTCGCTCAAATGCTTTCGCTCATCGCCTCGCATTTCTCACCTATTTCTC

GACCAGCGAAGATCAGAGGTTCGTGTTTCGAGCGGACAGTTCGGAACGGTTTCGGCCTTGGCGTCTGATCCTGATCAGTT
GAAGAGCGCTAGAGAAGATATCAAGGAGCTTCTCAGTACTAAGTTCTGTCATCCTCTTCTGATTCGTTTGGGATGGCATG
ATGCTGGTACTTATAATAAGAATATTGAGGAGTGGCCACAGAGAGGTGGAGCCAACGGTAGCTTAAGATTTGAAGCTGAG
TTGAAACATGGAGCCAATGCTGGACTTGTAAATGCATTGAAACTTCTCCAGCCGATCAAAGACAAGTATTCAGGCGTGAC
ATATGCAGACTTATTTCAGTTAGCCGGTGCTACAGCTGTGGAGGAAGCTGGAGGCCCCAAAATTCCCATGAAATATGGAA
GAGTAGATACCACTGGACCTGGCCAATGTCCTGAAGAAGGACGACTTCCTGATGCAGGCCCCCCTTCACCTGCTGATCAT
TTGCGTGAAGTTTTTTACCGAATGGGATTGGACGACAAGGAAATTGTTGCATTATCTGGTGCGCACACACTGGGGAGGTC
TAGACCAGACCGTAGTGGCTGGGGAAAACCTGAGACTAAATACACGAAAGACGGGCCAGGAGCACCTGGAGGACAATCCT
GGACTGCACAATGGTTAAAGTTTGACAATTCTTACTTCAAGGACATCAAAGAAAAAAGGGATGAAGATCTATTGGTATTG
CCAACTGATGCAGCTCTTTTTGAAGATCCTTCCTTCAAGGTATTTGCTGAGAAATATGCTGAAGATCAGGAAGCATTCTT
TAAAGATTATGCTGAAGCACATGCCAAACTCAGCAACCTTGGAGCCAAATTTGATCCTCCAGAGGGAATTGTGATAGATG
GTTCTCCAATTGTAAAGGGAGAGAAGTTTGTAGCAGCCAAGTACTCCTCTGGAAAGAGAGAGCTGTCCGATGCAATGAGG
AAGAAGATACGAGCTGAATATGAGGCAGTTGGCGGAAGCCCAGATAAGGCTCTAAAGTCAAACTATTTCCTAAACATCAT
AATTGTTGTTGCAGTTTTGGCTATTTTGACATCCCTGCTTGGAAACTAA
>evgLocus_145163
ATGGCACAATATAATAGAGTTGATGCTGAGTATGTTAAGGAAATCAACAACACTCGTAGCGATCTTCGCTCTCTTATTTC
CAACAAAAAATGTGCTCCTATCATGCTTCGTTTAGCATGGCACGATGCTGGTACCTACGATGCAAAAACAAGAACAGGAG
GTCCAAATAGTTCTATTCGAAATCAACAAGAGCTCAATCATTCTGCTAACAAAGGCTTGAAAATCGCCGTTGAATTACTT
GAGGAAGTGAAAGCTAAACACCCCAAAATTTCATATGCGGATCTTTACCAGCTAGCTGGTGTTGTGGCTGTAGAGGTTAC
TGGTGGTCCAGCTATTCACTTTGTTCCTGGTAGAAAGGATTTATTAGAATCTCCAGAAGAAGGTCGTCTTCCTGATGCTA
AACAAGGTGCATGGCATTTAAGAGAGGTTTTTTATCGCATGGGTCTCAATGATAGAGACATTGTAGTTCTTTCCGGAGGC
CACACTTTGGGTAAGGCACATAAGGATCTATCTGACTTTGAAGGTCAATGGACAAGAGACCCTCTTAAATTTGATAATTC
TTATTTTGTAGAATTATTGAATTCGGAATCAAAAGACTTTTTGAAGCTTCCCACTGATAAGGTTCTAGTTGAAGATCCTG
CATTTCGAAAATATGTTGAACTCTATGCTAAGGATGAGAAAGCTTTTTTCAGAGATTATGCAAAGTCACACAAGAAACTC
TCTGAACTAGGCTTTAATCCCAATGGCAATTATCTTTCCATATTGACTAAGGCAGTCTTAGGAGTGGTTATTGCATCAAC
TGTTGTGGTTCTGGGTTACTTGCTTGAACTCAACAAAAAAATCAACTGA
>evgLocus_145931
ATGGCGTTGCCGGTGGTGGTTGACGCAGAATACCTCAAGCAGATCGAAAAAGCTCGCCGCGATCTCCGAGCACTCATCGC
CAACAGAAACTGCGCTCCTCTCATGCTCCGTTTAGCTTGGCACGATGCGGGAACTTACGACGCCATCTCGAAGACCGGTG
GACCTAACGGCTCTATCCGCAACGAGGAAGAGTATTCTCATGGCGCTAACAATGGTTTGAAGAAAGCTATTGATTTCTGT
GAGGAAGTGAAGGCAAAGCATCCTAGAATCTCATATGCAGACCTTTATCAGCTTGCCGGTGTTGTTGCAGTTGAGGTTAC
TGGGGGTCCCACAGTCAACTTTGTTCCCGGAAGAAGGGATTCAAAAATATGTACCAGAGACGGGCGGCTTCCCGATGCTA
AAAAAGGTGAGTCGCATCTCCGTGATATCTTTTATCGTATGGGCTTGACTGACAAGGATATTGTTGCGCTGTCTGGGGCA
CATACACTGGGAAGAGCACATCCAGAGAGATCAGGGTTTGATGGCGCTTGGACAGAGGACCCTCTGAAATTTGATAACTC
ATACTTTGAGATACTTTTGGAAGAAGATTCTGCAGGGCTGCTTAAACTTCCAACGGACAGGGCTTTAGTGGATGATCCTG
AATTTCGCCGTTATGTTGAGCTCTATGCGAAGGACGAGGATGCATTTTTTCGAGATTATGCTGAATCACATAAGAAACTT
TCAGAGCTTGGCTTTGTGCCAAGCTCAAAGGCGAATGCTCCTAAGGATGCGACCATTCTGGCACAAAGTGCTGTTGGAGT
TGTAGTTGCTGCTGCAGCGGTCATCCTCAGTTACTTGTATGAACATTCCAGAAGAGGGAAGTAG
>evgLocus_255817
ATGGCTCAATATAGAGTTGACGCAGAGTATGTTAAGGAAATCGACAACACTCGTAGCGATCTTCGTTCTCTTATTTCCAA
CAAAAAATGCGCTCCTCTCATGCTTCGTTTAGCATGGCATGATGCTGGTACCTACGATGCAATAACAAGAACAGGAGGTC
CAAATGGTTCTATCAGGAATCAACAAGAGCTCAATCATTCTGCTAACAAAGGCTTGAAAATCGCCGTTGAATTATGTGAG
GAAGTGAAAGCTAAACATCCCAAAATTTCATATGCGGATCTTTACCAGCTAGCTGGTCTTGTGGCGGTAGAGGTCACTGG
TGGTCCAGCTATTCACTTTGTTCCTGGTAGAAAGGATTCGTTGGAATCTCCAGAAGAAGGTCGTCTTCCAGATGCTAAAC
AAGGTGCATCGCATTTAAGACAGGTTTTTTATCGCATGGTTATGAATGATAGAGACATTGTAGCTTTGTCCGGAGGCCAC
ACTTTGGGTAATGCACATAAGATCGCTCTGACTTTAAAGGTCAATGGACAAGAGATCCTCTCAAGTTTGATAACTCTTAT
TTTCGGGAATTATTGA
>evgLocus_256923
ATGAGAGAACTTGTGCACAAATATGAAAGAGAAGACAAGAGAATGTTTGTGATAATATGTGGGATCTCGTTGGTATTGTT
CATTGTAATCCAATGGTATTCCAAGTCTAGAAAAAGCAAGAACTCACCTCCTTCTCCTCCAAAACTACCCATTATAGGAA
ATCTCCATCAACTTGGTAAATTCCCCCACCGCACATTTCTATCTCTAGCTAAAAAATATGGCCCTGTCATGCAAATTCAT
CTTGGTAGTGTTCCATGTCTCTTGATCTCATCCTCCGAAGCAGCACGTGAGGTGATGAGAACCCATGATCATCATTTTGC
AGACAGACCCCAAAAGAACAATTACAAGATACTTATATATGATTGCAAAGACGTTTCAACTGCTCCTTATGGAGATTACT
GGAGACAGTTAAGGAGCATCTCTGTCTTGCATCTTCTCAGTGCTAAAAGGGTTCAATCTCTTCGCTCCGTACGAGCCGAA
GAAATCAGTTTAATGATGGAGAAGATAAAGCATTCTTCTTCTACTTCTGAGCCGCTTAATTTAAGTCAACTCATCGCTTC
CACAGTAAATGATATTGTTTGTAGGGTTGCTTTGGGAAGAAAGTATAGTGGTGAAGAAGGAAAGGGATTTAAAAAGTTGT
TCAGGGAATTTACCAATTTACTCGGTATGTTTATTGTTGGAGACTATGTACCTTGGCTTGATTGGGTCACCCATGTCTCT
GGAACTTATGGAAGAGCTAGAAGAGTCGCCAAACGCTTTGATGATCTTTTGGAGGATGTTGTTGAAGATCATATCAAAAC
TCACAAACAAGCCAATCATGATCTGGGTGAAGAAAACCATAAAGATTTTGTTGATGTTTTGCTTTGGATCCAGAGGACTG
AAGCACTTGGCTTTCCTATTGACAGAATTGTCATAAAGGCTCTCTTATTGGACATGTTTATCGGAGGTACAGGCACGACG
GCTAGTTTGCTGGATTGGGAAATGTCAGAGCTGATAAAGAATCCGCGCATGATGAAGAAATTGAAAGAAGAAATAAAAAG
TGTTGCAAATGGTAAAACACACATAACAGAAGACGATTTGGTTAACATGAAATACTTAAAGGCAGTGGTTAAAGAAACAC
TACGATTACATCCTCCATCACCGCTACTAATCCCTCGAGTAAACAAAGAAGATACCGAATTCAATGGTTACCATATAAAA
GCCGGGACACAAGTTATTATAAACATATGGGCGATTGCAAGAGATCCTGCAAATTGGGATTCACCAGAAGAATTCAAGCC
AGAAAGATTCTTGGACAGTACAATAGATGTTAAGGGTAATGATTTCACATTGATTCCATTTGGATCAGGGAGAAGAGGTT
GCCCTGGGGTTGTGTATGCTTTGTCTGTCAATGAAATTATGTTGGCCAACATTGTGCATCAGTTTGACTGGGAAATACCT
GGAGGCTTTGAGACATTAGATATGGCTCCATCAACTGGCTTTCTTGCGCATAAAAATACTCCTATTATCGCTCTTGCAAC
TCCTACTAGTAACTAA

```
>evgLocus_297956
ATGACCTCAAACAAACCACAAAAGATCTTACACTATTGCATCACCTTTGTTCCCCTTCTCCACTTCCAAACACACACCTC
AAACAATTTGGTTGTAGTTGCAACAACAATGGGTGTCTCTTTCCTCTCAACCTCAACTCTTCCTTCTCTCCATCCACTAC
TTCGTTCTCATTATCCTAAAACAACTACAACCACACACCAACCTCGTTCGGTTTCCATTTGTTGTTCTAAAATCCAAACA
GATGTTACTGATGAAGAACAACAGTTCCTTTGGAAACGAAGAGACATTCTTAAATGCTTCGGGGTCACTATTGGTTTGGA
ATCGATAACAAACTCTGGATTGCTTGTTGGAACGGCTAATGCTGCTGACCTGATTGAACGCAGACAGCGTTCGGAGTTTC
AATCTGAAATTAAAGGAACTCTTTATAAGGCCATTAAGGGAAATCCCGATATTGTTCCATCCATACTAACACTGGCGATA
AATGATGCTTTAACTTATGATAAGGCAACGAAAACAGGTGGCCCAAATGGCTCTATACGGTTCAGCTCAGAAATAAGTAG
ACCTGAGAATAAGGGGTTTTCTGCAGCCTTGAATTTCATAGAGGAAGCAAAAAAAGAAATAGATTCATATTCGAAAGGTG
GACCGATTTCCTATGCAGATCTAATCCAGTATGCAGCACAAAGTGCAACAAAGGCTACATTTTTAGCTTCTGCGATTCGC
AAATGCGGTGGAAATGTGGAAAAAGGGAACTTGTTGTACACAGCATATGGATCAAACGGACAGTGGGGTTTGTTCGACCG
ACAATTTGGTAGGGCTGATACTCAAGAGCCGGATCCAGAGGGGAGAATTCCAATTTGGGAGAAAGCAAGTGTTCAGGAAA
TGAAAGATAAGTTTTCTGCTGTAGGCCTTGGTCCTCGCCAGCTTGCTGTTTTATCTGCATTCATAGGTCCGGACCAGGAT
GCAACAGAAGCCTTATTAGCATCTGATCCAGACGTTGCTCCATGGGTAACAAAATACCAACGGAGCCGTGAAACTGTCTC
GCGAACCGATTATGAGGTTGATCTTATAACAACTTTCACAAAATTGAGTACCTTGGGCCAAAATATTAACTATGAAGCAT
ACACGTATCCTCGTAAGAAGATCGACATTACTAAACTCAAGTTGTAA
>evgLocus_366734
ATGGATTATGCGATTAGTGCACTGCTCATTTTGTTGACATGCATTGTCACATACTTTGTTCGTTCACTTTCTGCAAGAAC
CAAAATTTCAGACTACAAGCTTCCACCAGGACCTTCCTTTTTCACTATCATGTCAAATGTTGTTGATTTGTACAACAAGC
CACAACAAACACTTGCAAAATTTGCTAAGTTCTATGGTCCTGTTATGCGTATAAACCTATGCACTGAAACCACTATAATA
ATCTCGTCGTCTGATATGGCCAAAGAAATTCTCCATACTCATGATTCTTTGTTCACTGATAGATCTGTTCCTCATAATAC
TACAATTTACAACCACAACAATTTTAGCTTAGTCTTCCTCCCATTTTCACCTCTTTGGCAACACCTTAGGAAAATATGTC
ATAATAATCTATTCTCTAGCAAGACCCTTGACGCAAGTCAAGAACTTAGACGAATTAAACTAAAAGATCTTCTCAATGAT
ATGCATAAAAGCAGTTTAACTGGTGAAGCTGTTGATATTGGAAGAGCTGCTTTCAAAGCTTGTATTAATTTTTTGTCGTA
CACTTTTGTGTCTCAAGATTTTGTTGACTCTCTGGATGATGAGCATAAGGATATAGTTTCCACTCTTCTGAAAGCCATTG
GAACACCAAACATCTCTGATCATTTCCCTGTGTTGAAGATATTTGACCCACAAGGGATCAAAAAACTCACTTATAATTAT
GTTTCAAAGGTTTTTTTTGCCTTAGATATAATAATTGAGAAGCGAATGAAGTTGAGGGAAAGTGAACATCATATCTCAAA
CAATGACATGTTAGACACTTTGTTGGACATTTCCAAAGAAGATAATCACAAGATGGATAATAAGCAGATTAAACATCTAT
TACTTGATTTGCTTGTGGCGGGAACAGATACAACAGCATACGGATTAGAAAGAGCACTGAGTGAACTAGTACACAATCCA
GAGATTATGTCAAAAGCCAAAAAGGAACTTGAGGAGATTATTGGATTAGGAAATCCAGTTGATGAATCAGACATCGATAG
GCTTCCATACTTACAAGCAGTGGTAAAAGAGAGTTTACGTTTACATCCTCCAGCTCCAATGTTGCTCCCTCGAAAAGCAA
GGGTAGATGTGGAAATATCAGGATACACAATTCCAAAGGGTGCTCAAGTTTTGATCAATGAATGGGCTATTGGTAGAACA
GACATATGGGAAGATGCTCATTTGTTTTCACCAGAAAGGTTTATAGGATCTGAAATAGATGTAAAAGGTCGACACTTTAA
GCTTACACCATTTGGCAGTGGGAGACGTATATGTCCAGGATCACCATTGGCAGTGAGGATGTTGCATTTGATGTTGGGAT
CATTGATCAACTCATTTGATTGGAAACTTGAAAACAATATGGAGTTTAAAGACATGGATTTGGACAATTCTCTAAGAGCT
ATTCCGGTTGCACTAAACAAAGTGTTATAA
>evgLocus_727341
ATGACTCTGTTTCTCACCATACCTCTTTCACTTCTCACCCTCTTCATCTTCTACACCCTCTTCCAACGTCTCAGATTCAA
GCTTCCACCCGGTCCACGCCCCTGGCCGGTCGTCGGAAACCTCTATGACATAAAACCAGTCAGGTTCCGGTGTTTCGCGG
AATGGACCCAGTTCTACGGGCCAATTATATCGGTTTGGTTCGGTTCGACTTTGAACGTGATTGTTTCGAATACGGAGTTG
GCGAAAGAGGTTTTGAAAGAGAATGATCAGCAGTTGGCGGACCGGCATAGGAGTCGGTCGGCTGCCAAGTTTAGTAGAGA
TGGGAAGGATTTGATTTGGGCTGATTATGGACCTCATTATGTGAAGGTGAGAAAGGTTTGTACTTTGGAGCTTTTTTCGC
CCAAGAGAATTGAAGCTTTGAGGCCTATTAGAGAAGATGAAGTTACTGCTATGGTTGAATCTATTTTTAATGATTCTACT
AATCCTGAAAATTTGGGGAAAGCTATACCAATGAGGAAGTATATAGGGGCGGTTGCATTCAACAACATCACAAGGCTGGC
TTTTGGGAAAAGATTTGTGAACGCAGAAGGTGTAATGGATGAGCAAGGAGTAGAATTCAAGGCTATAGTGGCAAATGGGT
TAAAGCTAGGAGCATCTCTAGCTATGGCAGAGCACATCCCTTGGTTGCGCTGGATGTTCCCACTAGAAGAGGAGGCTTTT
GCCAAGCACGGTGCTCGCCGAGACCGACTTACTCGAGCCATCATGGACGAGCATACACAAGCACGCCAGAAATCCGGCGG
TGCTAAGCAACATTTTGTAGATGCCCTTCTCACCTTGCAAGACAAATATGATCTTAGTGAAGACACCATCATTGGTCTCC
TTTGGGACATGATAACTGCTGGGATGGACACAACTGCAATATCGGTTGAGTGGGCCATGGCCGAGTTGATAAAGAATCCA
AGAGTGCAACAGAAGGCACAAGAGGAGCTAGACAAGGTCATTGGTTTTGAAAGGGTCATGACAGAAACTGACTTTTCAAG
CCTTCCTTATCTACAAAGTGTAGCCAAGGAGGCTCTAAGGCTGCATCCACCAACACCATTAATGCTCCCACATCGCGCTA
ATGCGAATGTTAAAATTGGTGGCTATGATATTCCCAAGGGGTCCAATGTCCATGTCAATGTATGGGCGGTAGCTCGTGAC
CCGGCTGTATGGAAAAACCCGTTGGAGTTTAGGCCCGAGAGGTTTCTTGAAGAGGATGTAGACATGAAGGGACATGATTT
TAGACTACTTCCATTTGGAGCAGGTCGTCGGGTTTGCCCAGGTGCACAACTTGGAATCAATATGGTGACATCGATGTTGG
GTCATCTATTGCATCATTTCTGTTGGGCACCTCCTGAGGGAGTGAACCCTGAGGAGATTGATATGGTAGAGAACCCTGGA
ATGGTGACATATATGAGGACTCCATTACAGGTTGTGGCTTCTCCAAGGCTTCCCTCAGATTTATACAAACATGTGCCAGC
TGATATCTAA
>evgLocus_959121
ATGGGAAAATCTTACCCTACCGTTAGTCCTGATTACCAGAAGGCCATTGAAAAGGCCAAGAGGAAGCTTAGAGGTTTCAT
TGCTGAGAAGAAATGCGCTCCTTTAATTCTCCGTTTGGCATGGCACTCTGCTGGTACTTTTGATTCGAAGACAAAGACTG
GTGGTCCTTTCGGAACCATTAAGCATCCAGCTGAGCTTGCTCATGGTGCTAACAACGGTCTTGATATTGCTGTCAGGCTT
TTGGAGCCTCTTAAGGAGCAATTCCCTATTGTGAGCTATGCTGATTTCTATCAGTTGGGTGGTGTTGTTGCTGTTGAGAT
CACTGGTGGACCTGAAGTCCCTTTCCACCCCGGACGGGAGGACAAGCCCGAGTCACCACCAGAGGGTCGTTTGCCTGATG
CCACCAAGGGTTCTGACCATTTGAGGGATGTGTTTGGAAAAGCTATGGGGCTTAGCGATCAGGACATTGTTGCTCTATCT
GGTGGTCACACCATTGGAGCTGCACACAAGGAGCGTTCTGGATTTGAGGGGCCATGGACTTCTAATCCTCTCATTTTTGA
CAACTCATACTTCACTGAGTTGTTGACTGGTGAGAAGGAAGGCCTTCTTCAGTTGCCAAGTGATAAGGCACTTTTGTCTG
ACTCTGTATTCCGCCCTCTTGTTGAGAAATATGCTGCGGATGAAGATGCCTTCTTTGCTGATTATGCCGAAGCACATCTT
AAGCTCTCCGAGCTTGGATTTGCCGAAGCCTAA
```

```
>evgLocus_1209062
ATGTGTTCAAGTGCGAGTAACCGTATTGTGCAGTGCTCAGTCGCAACCGCACCAGGTGGTGAAACTAATTCTCCCTCCAA
ATTACAACCACCGTCTTTCTCAACCGTAAACTTTCGCTCCGTGAGACCCAACGACCATGTTTCTTCCGAGGTTGCGAGTA
GCAGCAGGAGAGGGCTAATATGCAGCATAGCCATGTTGCCCTGTCTTCTTCCACTCACTCACATCTCTGGTTCTCTCCCA
GCCAACGCCATGCCACTACCAGATACAGAAGAATATGTTCGAATTAAACAAGAGTTGAGGAAGGTATTGTCAAAGGGAAA
GGCTGCAGGCGCGCTTCGTTTGGTTTTTCATGATGCTGGAACTTTTGAAATTGATGACAATACAGGTGGCATGAATGGCT
CTATAGTCTACGAACTTGAAAGACCTGAAAATGCCGGTCTGACAAAATCAGTGAAGGTTCTGCAGAAAGCCAAGACTCAG
ATAGATGCAATCTATCCAGTATCCTGGGCGGACGTTATTGCTGTGGCTGGAGCTGAAGCAATTGAACTATGTGGAGGTCC
TACTATCCAAGTTTCACTCGGCAGACAAGATTCGCTGGGGCCTGATCCTGAAGGAAAACTTCCTGAAGAAACTCTTGATG
CTTCTGGTTTGAAAAGATGCTTCAAGAAAAAAGGCTTTTCAACACAAGAACTGGTAGCTTTGTCTGGAGCTCACACTCTT
GGAAGTAAAGGTTTTGGAAGCCCTACATCTTTTGACAATTCATATTATAAGGTTCTGTTGGAAAATCCACCCACGGCTTC
TGGCGGTTTGTCGACTATGGTTGGTCTTCCTTCAGATCATGCACTTGTTGAGGATGAAGAATGCCTCAGATGGATTAAAA
AATACGCAGACAATGAGAATATGTTCTTTGAAGATTTCAAAAATGCGTATGTCAAGCTGGTGAATTCTGGTGTGAGGGTC
ATTAAATTTGTGTACCAAAGTGTTTAA
>evgLocus_1273529
ATGGCATTTTCAACACATGAGCAACAGCATCAACTATTATCAAACCTTCACGGAGGTGCCACGTCAGCACCGCCACCAAC
TCCGTCAAATAACACCAACAACCTCCTCTCATCTCCCGACGCCGCCGACGCACTCTCCCGCCTTCTCCACCGTCTCCCGC
CTAATCTCTCTCTCCCAAACCGCCGCTCCTCTTCATCCGCCACGTCTCCTCCGTCTCTCTCCTTTTCTTCATTAACTCCA
AACGAACTTCTCTCTTCTGTTTCCAAACTCGGTTTTATCCAACTCAATGATCACTCAGTTTCATCTGAACTCGCCAAATC
GGCCGAGTCGGAATCGCTTAAACTATTCGACCTTTCTCGTGACCAGAAGGAAATTTTCTTCCCTCAAAACTGGCCGTTAG
GTTACGAAGGCGGCGCCAATGACGACGGTGACGGAATCTCTGAGTCGTTCCGGTTAGACTTACCGTGTTCAACCGAGTCA
AACGACTTAAAACTAGATTCAGTGAACGAGTTCGCACGCGCGCTTGAGAAGGTGGGTTTGAATATCATTGATGTGTTAAC
GAAGGGTTTAGGAGTTGTGAATCCGGCTGAAGAGGACCGAACCCGGTTCAGTTCGATTATGTGGATTTCGGAGGGAAATA
AACCCGGTTCAACGAGTGGGTTTTACCCGTTTATCATTGGGTTGCAGTATCAGATAAGGTGCCAGAAGCACTCTTTGTTG
TCTGATTCAAGTGGATGGGTTTCTGTTTTGCCACACGTGGACTCCATCTTAGTTACTGTTGGTGATATTGCCCAGGTTTG
GAGCAATGGAAAGCTAAAGAAAGTGAGAGGCAGACCAGTGGCAACATTGGGAGATGAAAATGATTCACGTTGCATAACAA
TGTCATTGTTGATAACTCTTCCTACAGAGAGCAATGTAGCTCCACTTCTTCCCATTGGCTTCAAAGACAAAGTTGAAGAT
GATGAAGAAGAAGAAGAGAGCAACATTGTTGGTGAAGTCCAAAAGAGGGTATTCAACTCCATTGATTTTGAGGACTATGC
TTGGAGAGTTTATCATGAAAATCTCTTATTCAAGGATCCATTGGATAGGTACCGTGTGACTCAATAG
>evgLocus_1282817
ATGCTAGCGATAATCTGTTCCAGTTTTGTGTTTCTTTTTCTTGTAATAAAATTGTATTCCTATTCTTCAGAAACCAAAAA
CTCACCTCCAAGATTACCTTTAATAGGAAATCTTCATCAACTTGGTTCATTCCCCCACCGTTCTTTTTACGCTCTAGCCA
AAAAATATGGTCCTTTCATGCAAATGTATTTTGGTAAAGTTCCTATTCTTGTCATCTCATCAGCTGAAGCAGCACGTGAG
ATAACCAAAACTCATGATCATGTCTTTGCAAACAGACCCCCCAAGATTAATTACGACATTCTTTTGTATAATTTCAGAGA
TGTTTCATCTGCTCCATATGGAGAGTATTGGAGACAGTTAAGAAGCATTTGTATGTTGCATCTTCTCAGTGCCAAAAGGG
TTAAATCTCTTCGCACTGTGAGAGAGGAAGAACTTGTTTTTATGATGGATAAAATAAGAGACTATTCTTCTAAATCACTG
CCTGTGAATTTAAGCGAATTGATTGCGTCAAAAACTAATGATGTAGTTTGTAGGGCTACTTTGGGAAATAAGTATAGTGG
TGAAAGTGGAACAGGATTTGCTAAGTTGATGTTGGATTTCACCGAGTTGCTTGGTACTTTTATGGTTGGGGATTATGTTC
CTAGCCTTGATTGGATGACACATCTTTCTGGATATTACTCCAGAGCAAAGAAAGTCGCCAAACAATTTGACGATCTTTTG
GAGGGTGTAGTCGAGGAACGTTTCAATAATCCGAAAGGTGATGATGAAGAACAGACTGATTTGGTTGATGTTTTGCTTTG
GATCCAAAGGACTGAATCCCTCGGCTTTCCTATCGATAGAACAACCATAAAGGCTTTGTTACTGGACATGTTTGTTGCGG
GTACAGACACCATATCAACTTTGCTAGAGTGGGAAATGTCAGAACTGTTGAAGAATCCACACATGATGAAGAGATTAAAA
GAAGAAGCAAGGACAGTGGCTAACGGAAGAGCATACATAACGGAAGACGATTTGAGTAACATGAAATACTTAAAGGCACT
TGTTAAAGAAGCACTACGGATGTATCCTCCGATCCCACTACTAGTCCCTCGAGAATGCAGACAAGATGTGAAAGTAGACG
GTTACAACATTAAAGCCGGGACAAGAGTTTTTATCAACGCTTGGGGAATTGCAAGAGATCCACGATATTGGGATCAACCT
GATGAGTTCAGGCCAGAGAGATTCTTGGATACTTCGGTGGATGTTAAAGGAATTGATTACCAGTTGATTCCGTTTGGATC
GGGGAGAAGAGGTTGTCCTGGACTAGTGTATGCTATGGCTGCTAATGATCTTGTGTTGGCCAACCTTGTGCATCAGTTTA
ACTGGGAGTTACCTGGTGGTGCTGGGGCCAAGGTCGATATGTCTGAAGCATTTGGTTTTACTGTCCACAGGAAGTTTCCT
CTTATGGCATATGCAGCCAGCACTTTCTAA
```
